# Time-resolved genome-scale profiling reveals a causal expression network

**DOI:** 10.1101/619577

**Authors:** Sean R. Hackett, Edward A. Baltz, Marc Coram, Bernd J. Wranik, Griffin Kim, Adam Baker, Minjie Fan, David G. Hendrickson, Marc Berndl, R. Scott McIsaac

## Abstract

We present an approach for inferring genome-wide regulatory causality and demonstrate its application on a yeast dataset constructed by independently inducing hundreds of transcription factors and measuring timecourses of the resulting gene expression responses. We discuss the regulatory cascades in detail for a single transcription factor, Aft1; however, we have 201 TF induction timecourses that include >100,000 signal-containing dynamic responses. From a single TF induction timecourse we can often discriminate the direct from the indirect effects of the induced TF. Across our entire dataset, however, we find that the majority of expression changes are indirectly driven by unknown regulators. By integrating all timecourses into a single whole-cell transcriptional model, potential regulators of each gene can be predicted without incorporating prior information. In doing so, the indirect effects of a TF are understood as a series of direct regulatory predictions that capture how regulation propagates over time to create a causal regulatory network. This approach, which we call CANDID (*Causal Attribution Networks Driven by Induction Dynamics*), resulted in the prediction of multiple transcriptional regulators that were validated experimentally.

## Introduction

A central problem in modern genomics is how to extract causality from experimental data. Additionally, distinguishing direct from indirect effects is an enduring challenge. In experiments that elicit dynamics (often due to environmental perturbations), linking responses to potential upstream molecular causes to build gene regulatory networks (GRNs) can be done with the aid of prior knowledge [1]. Integrating prior knowledge with genomic studies of mutants has also been used to determine *direct* regulatory relationships between transcription factors (TFs) and genes involved in core cellular processes, from cell cycle control to the DNA damage response [2,3]. GRNs typically focus on well-studied genes, require extensive prior information to elucidate, and are often missing direct molecular interactions. Addressing these issues requires a fresh perspective.

Large-scale maps have been generated with the goal of identifying direct TF regulatory targets [4,5]. Interpreting the biological impact of these interactions is challenging because regulatory interactions are dynamic and contingent on physiological state. Genes with similar ChIP profiles can exhibit opposite expression responses [6], and highly expressed regions of the genome can be “hyper-ChIPable”, resulting in a non-biological source of signal [7]. Alternatively, genetic perturbations with expression or growth-rate as readouts can be used to group functionally similar genes and processes [8,9]. Gene expression profiling of deletion mutants (i.e., asymptotic readouts of a perturbation) can help identify co-regulated genes, though without dynamics there is limited potential for determining direct regulatory relationships because how information propagates from the deletion to each differentially expressed gene is not observed [10].

We argue that identifying *direct causal* relationships without prior knowledge can be improved by using well-defined interventions followed by longitudinal genome-scale measurements. The work of McIsaac *et al.* adopted this approach for the purposes of dissecting the incompletely understood regulatory connectivity of the yeast sulfur regulon [11]. By generating strains that expressed each known sulfur-related TF from an engineered promoter that could be activated by a small molecule (β-estradiol), a single TF could be rapidly induced and responses could be tracked as they propagated over time. But genome-wide time-resolved datasets are uncommon, and accordingly, existing approaches utilizing dynamics must either rely extensively on prior information to predict the true regulator among a set of correlated alternatives [12], or focus on small networks where many possible regulators are removed [1,13]. Thus, the field requires experimental datasets that are suitable for elucidating GRNs, and new analytical approaches for learning non-canonical regulators from such data.

Here, we present an approach (CANDID) for revealing genome-wide causal relationships without incorporating prior information. We generated over two-hundred TF induction timecourses in which a single yeast TF was rapidly induced, and full transcriptome differential expression was tracked, typically across eight timepoints. Such timecourses feature the near immediate strong induction of an inducer-driven TF of interest, followed by rapid changes in genes that are directly regulated by these TFs, and later changes of indirectly regulated genes (Figure 1A-C). While these indirect effects contain many uncharacterized regulatory processes, they can be difficult to attribute to a single specific regulator (Figure 1D) using single timecourses. By aggregating all timecourses, we can more confidently identify which regulator(s) are acting in each individual timecourse by finding the parsimonious set of regulators whose abundances account for each gene’s expression variability (Figure 1E, F). Furthermore, our approach implicitly dissects indirect regulation into a series of direct regulatory relationships, and by not utilizing prior information, we minimize bias against re-learning known biology. Accordingly, predicted intermediate regulators span canonical transcriptional regulators as well as genes of unknown function. We tested ten predicted latent regulators, and found three of them to be genuine transcriptional regulators and are commonly induced and repressed across many experiments.

**Figure 1.**
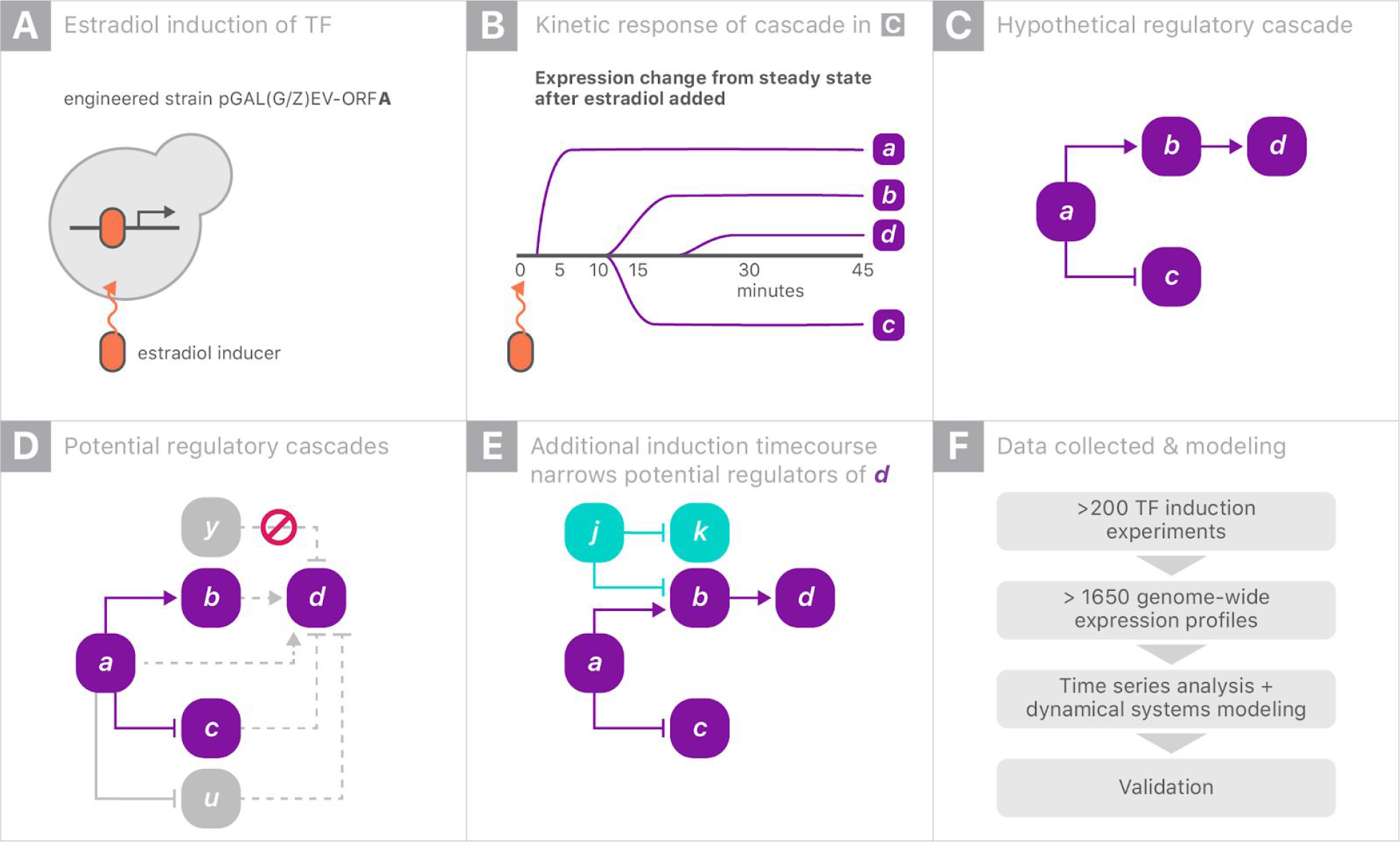
Inferring direct regulation using overlapping induction timecourses. **A.** In each experiment, one transcriptional regulator with an inducible promoter is rapidly overexpressed in response to 1 μM β-estradiol. **B.** Example of three genes (labeled *B*, *C*, and *D*) responding with different kinetics following induction of regulator *A*. **C.** Hypothetical example of a regulatory cascade in which an induced transcriptional regulator *A* directly inhibits *C* and directly activates *B*. *B*, in turn, directly activates *D*. **D.** In practice, we don’t know that *A* regulates *D* via *B* and instead want to infer such regulatory relationships. In this example, direct regulation of *D* by *B* is only one hypothesis that is consistent with the data - all viable hypotheses are shown by dashed lines. *A* could directly activate *D*, *C* could inhibit *D*, or *A* could regulate an unmeasured confounder *U* which is the true regulator. Direct regulation by a variable *Y*, which is independent of *A*, is not possible since the timecourse begins at steady state. **E.** Integrating the *A* induction timecourse with a second induction timecourse, which perturbs *B* without perturbing *A* or *C*, allows us to narrow down *D*’s possible sources of regulation. In this case, *U* may still be a possibility if it is remains correlated with *B*. **F.** Overview of data and analysis performed in this study. Over 200 induction timecourses were constructed allowing for many opportunities to resolve ambiguous regulation.

## Results

Each of 201 genes’ native promoters were separately replaced with a β-estradiol-inducible promoter as previously described [11,14] (Table S1). This set of induced genes is heavily enriched for non-essential TFs and chromatin modifiers. Each strain was grown to a steady-state in chemostat culture and following the addition of β-estradiol, the full transcriptome was measured at 4-10 post-induction timepoints (83% of timecourses contained 8 timepoints). Initially, 1691 microarrays were generated from 217 distinct induction experiments. 15 induction experiments were repeated at least once using the same induced gene to either capture late changes in some experiments (e.g., *GCN4*) or to allow for calibration experiments across the two induction systems used in this study (referred to as ZEV [14,15] and GEV [16]).

Most genes’ expression did not change in a typical induction experiment, with some notable exceptions (i.e., induction of Gcn4). Accordingly, the inducer-driven signal of interest is relatively sparse and interspersed among ubiquitous noise. This noise was governed by both a mild stress response [17] and Gaussian noise that varied across both genes and arrays. In order to isolate inducer-specific expression changes, the stress response was subtracted from each timecourse and then an observation-level noise model was used to select a subset of timecourses that are statistically inconsistent with experimental noise (Figures S1, S2, S3). The signals from these 100,036 timecourses were retained (~8% of timecourses) while all other timecourses were set as invariant (a log2 fold-change of zero). The full transcriptional dataset is available as Table S2. Further details on processing these data can be found in the supplement.

### Regulatory cascades and impulses

The Aft1 timecourse is an illustrative example of the value of induction data for revealing intricate regulatory phenomena. When Aft1 is induced, two broad classes of expression changes are observed: fast induction of targets which are known to be bound by Aft1 based on ChIP and gradual changes of genes whose expression has previously been shown to be correlated with, but not bound by Aft1 (Figure 2A) [18]. Such expression changes were typical of our dataset. Most genes in this dataset exhibit either a sigmoidal or impulse response (double sigmoidal); thus, we fit a Bayesian version of the Chechik & Koller kinetic model to each timecourse [19]. These parametric fits enabled the direct comparison of timecourses based upon whether they were sigmoidal or impulses and by using interpretable kinetic parameters. Sigmoidal responses are summarized with a half-max time constant *t*_*rise*_ and asymptotic expression level *v*_*inter*_. Impulse responses include two additional parameters: t_fall_, which describes the time when the response returns halfway to its final level, and *v*_*final*_, the asymptotic expression level of the impulse (Figure 2B) [19]. Utilizing these kinetic parameters, we observed multiple binding motifs of genes with characteristic response kinetics, including, as expected, the Aft1 motif associated with early activation, and a different motif (recognized by Tod6/Dot6 [also referred to as a PAC motif]) associated with early inhibition (Figure 2C). Targets activated and repressed in the Aft1 experiment have similar kinetic responses, and both classes contain examples of impulse-like expression responses (Figure 2D). Beyond Aft1, other TFs with a large number of impulse-like responses (indicative of feedback control / perfect adaptation) include Pho4, Mac1, Oaf1, Rtg1, Rtg2, Stb5, and Zap1 (S4, S5, S6). In total, we find evidence of transcriptional feedback in more than 1700 timecourses (~2% of all timecourses). To allow others to provide comparable investigations into the kinetics, functional coherence, and regulation of each timecourse in our dataset, we provide an interactive website (http://candid.research.calicolabs.com).

**Figure 2.**
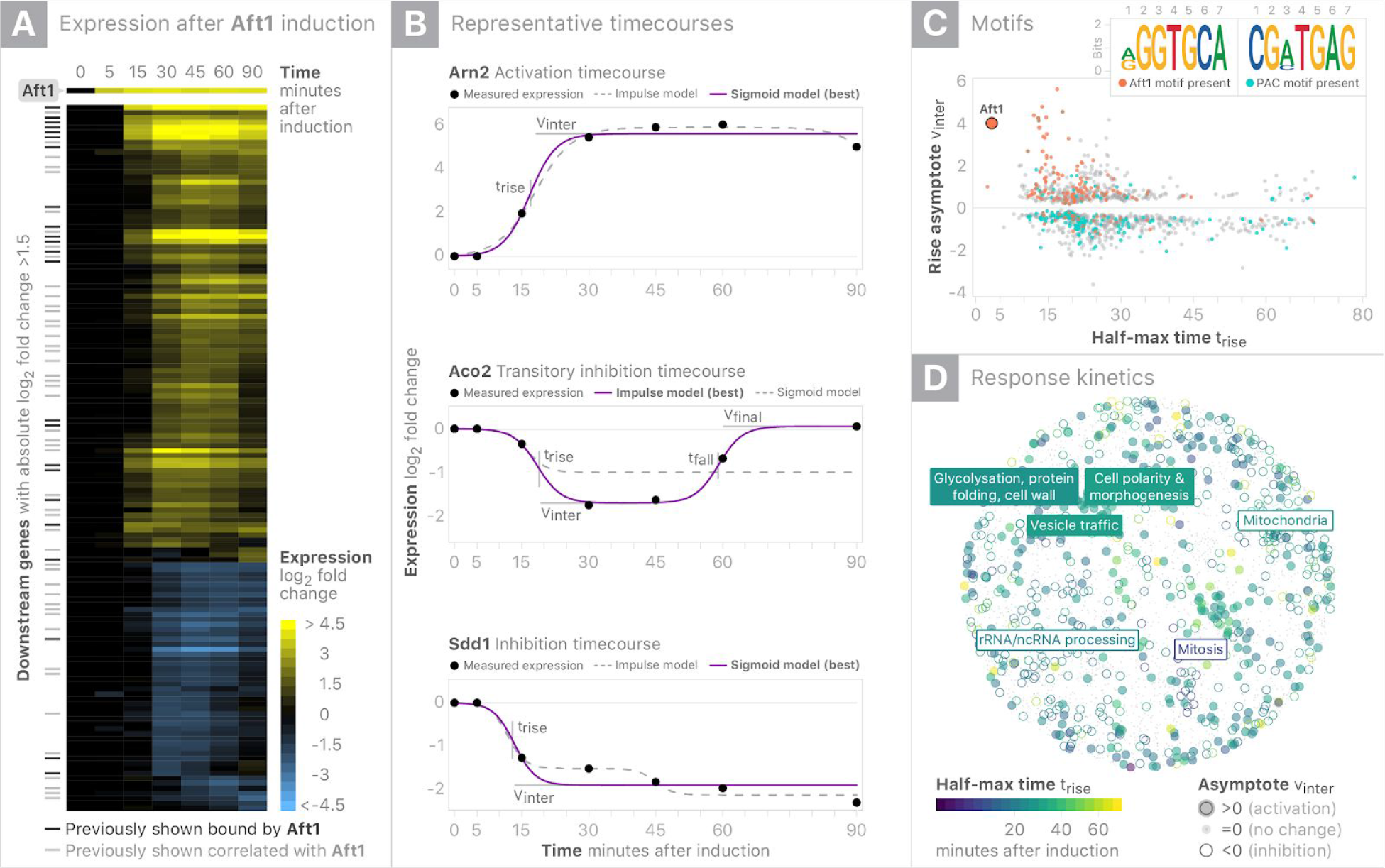
Characterizing the downstream responses of Aft1 induction. **A.** Heatmap summary of Aft1 induction timecourse showing all genes that change with |log2(fold-change)| > 1.5. **B.** Parametric summaries of representative sigmoidal activation and inhibition timecourses and impulse (double sigmoid) modeling of transitory inhibition. Sigmoids are summarized by half-max time (t_rise_) and the asymptote (v_inter_), while impulses include a second half-max (t_fall_) time and final asymptote (v_final_). The strongest supported model for each timecourse is shown as a filled in line, while the alternative model is shown with a dashed line. **C.** K-mers enriched in the promoters of regulated genes are overlaid on summary of each gene’s t_rise_ and v_inter_. Presence of the Aft1 motif is associated with early activation, while early inhibition is associated with the PAC motif (Tod6/Dot6). **D.** Response kinetics are overlaid on gene coordinates based on synthetic lethality as a surrogate for functional similarity. Up-regulated genes are enriched for vesicle trafficking/glycosylation processes and down-regulated genes are enriched for mitochondrial/mitotic/rRNA processes.

We can broadly categorize timecourses at a dataset-level based on existing knowledge. While strong acute regulation events are frequently associated with the direct binding of the induced TF, over 75% of genes responding in our dataset are new regulatory connections (Figure S7). Additionally, we find that 79% of genes reported as being directly bound by a TF do not exhibit a significant expression response in the corresponding TF’s induction experiment (Figure S7). The low recall of reported transcriptional regulation underscores the value of dynamic data for evaluating realized regulatory potential. Actual regulation may be greatly impacted by chromatin accessibility and the regulatory context of the extracellular environment [11,20–22]. This is further supported by the weak agreement between the reported binding and coexpression partners of a TF with the number of genes that change when it is induced (Figure S8).

Since induced TFs directly account for a small portion of the observed expression changes, it raises a broader question: *which regulators are actually acting in each induction experiment?* To investigate whether the kinetics of responding genes can be informed by promoter composition, we carried out systematic *de novo* motif discovery of all timecourses and identified 715 promoter motifs enriched in the responding genes across all experiments (Table S3). 34% of these motifs could be matched to known regulators and thus suggest plausible candidates for regulators which may operate in each timecourse. While linking TFs to their targets using motifs has been a common assumption in order to enable genome-scale GRN inference, we find this assumption can be limiting. Indeed, in the Aft1 induction experiment, the Aft1 motif is associated with direct activation of only a small number of genes. Since we would also like to understand regulatory cascades without requiring regulators to possess direct DNA binding ability, we developed a model that, assuming no prior information, could allow for the elucidation of regulators with unappreciated transcriptional impacts.

### Integrative modeling

In a given TF induction experiment, we infer which early-responding genes are causally responsible for the gene expression changes that occur later in the experiment. In a single timecourse, however, we would only be able to identify a coexpressed cluster of genes whose expression coincides with a late change, rather than a single candidate regulator (Figure 1D). While reliably inferring regulatory mediators from a single timecourse is a dubious prospect, across all timecourses, genes respond in a median of twelve induction experiments (or in ~5% of experiments; Figure S9). Therefore, aggregating multiple experiments provides the potential to decouple each gene’s expression dynamics from those of spurious correlates. As we have generated hundreds of strong orthogonal gene-level perturbations, our dataset provides an opportunity to test this approach. Across >1650 samples, each gene has a distinct pattern of variation and establishing such expression distinctness required a dataset on this scale (Figure S10).

To learn direct regulatory relationships from such data, we formulate a set of gene-level regression models that attempt to predict the rate of change of each target gene as a sparse linear combination of all genes’ expression:

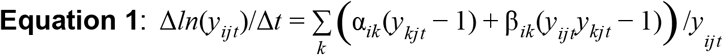

Here, *y*_*ijt*_ is the expression relative to the control strain and relative to time zero of a gene *i* in a timecourse *j* at a time *t* (i.e., for treatment *r* and control *g*, *y*_*ijt*_ = (*r*_*ijt*_/*g*_*ijt*_)/(*r*_*ij0*_/*g*_*ij0*_); therefore, *y*_*ij0*_ = 1 ∀{*I*, *J*}). Here, α represents the linear effect of one transcript on another and β represents the effect proportional to the target transcript. We allow any transcript to affect any other transcript, and thus we sum over all genes (with index *k*). Since most genes will not be regulatory, we use L1 regularization (i.e., LASSO) to shrink uninformative predictive coefficients to zero. We also enforce a predicted rate of change of zero at time zero, reflecting the pre-induction steady-state assumption. To arrive at this formula, we considered a suite of alternative data cleaning and modeling approaches (see Supplement for details) and decided upon this formalism and hyperparameters based on an ability to predict held-out induction datasets (in total, 50 million regressions performed).

The global regression model attempts to predict the rate of change of a target gene based on other regulators (Figure 3A). These instantaneous estimates can in turn be integrated to provide the model’s estimates of log_2_ fold-changes (Figure 3B). Grossly, the above model explains 43% of the variability in log_2_ fold-changes (Figure S11). While the model appropriately does not account for all expression variability, the variables in the above regression are directly determined by experimental data. Accordingly, our inability to predict one gene’s regulation does not affect modeling of another gene’s regulation.

**Figure 3.**
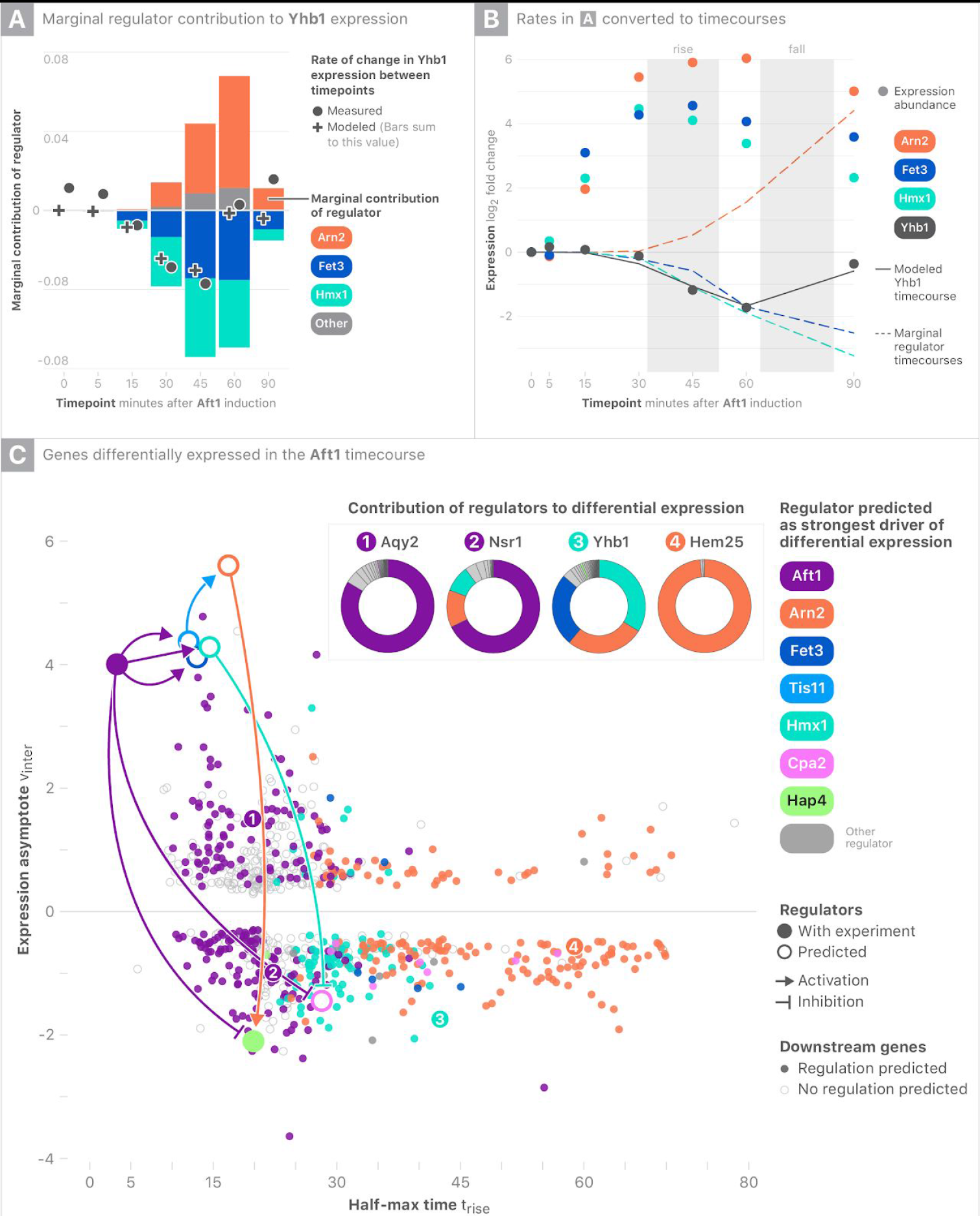
Predicted causal attribution of Aft1-driven transcriptional changes. **A.** Using a representative Aft1 target gene, *YHB1*, fold-change differences between timepoints (solid grey dots) are compared to the LASSO regression model’s fit (crosses). The model’s predicted marginal contribution of three predicted regulators (Fet3, Hmx1, and Arn2) to the combinatorial control of *YHB1* are shown with bars that sum to the model’s overall fit (crosses). **B.** The *YHB1* fold-change differences fit by regression can be converted to full timecourses to determine the marginal contributions of each regulator in driving a regulatory transition of interest (rises and falls). **C.** Each differentially-expressed gene in the Aft1 timecourse is laid out based on its kinetics and colored according to the regulator predicted to be the strongest driver of differential expression. Model-derived fractional contributions of regulators to expression of *AQY2, NSR1*, *YHB1*, and *HEM25* are depicted as donut charts.

### Decomposing indirect regulation into sequential direct regulation

The parameters of Equation 1 are regression coefficients that approximate ∂*y*_*i*_ / ∂*y*_*j*_; in other words, they capture the potential of a gene *j* to alter the expression of a gene *i*. However, to understand regulatory phenomena like the impulses observed in Aft1, we must consider both regulatory potential, as well as each regulator’s realized expression. To capture such relationships, we interrogate the fitted timecourses from our model, and to attribute changes to individual regulators, we consider their marginal contributions to overall timecourse changes. Using this framework, we look at each differentially expressed gene in a given timecourse and attribute each regulatory response (e.g., rise, or fall) to one or more regulators based on regulators’ marginal contributions to the response. Revisiting the Aft1 timecourse, our marginal attribution analysis predicts that different regulators are responsible for genes that respond with different kinetics (Figure 3C). In line with binding data, Aft1 is predicted to be the primary regulator of early activated genes, while Aft1 is predicted to turn off genes (in part) through the activation of Hmx1. Each regulator-target relationship can be thought of as a directed edge in a graph, with the whole graph describing how the regulation is predicted to have unfolded across time during each induction experiment. Performing such attribution analysis for all timecourses indicates that the induced TF is the primary direct driver of gene expression changes in nearly every experiment with signal (Figure S12); however, numerous other regulators are predicted as mediators of indirect effects.

The synthesis of timecourse-level graphs across all induction experiments reveals a global Causal Attribution Network (Figure 4) that links regulators with induction experiments to predicted intermediate regulators and the biological processes targeted by each regulator. In some cases (e.g., the Pho4 induction experiment), regulators predicted by the model do not connect back to the induced TF. This highlights that some regulatory phenomena are not explained by the model, but by directly using each gene’s abundance, its potential for regulation can be established. This global view reveals several predicted regulatory hubs, which are predicted to be directly activated or inhibited by multiple regulators and subsequently regulate a set of downstream targets.

**Figure 4.**
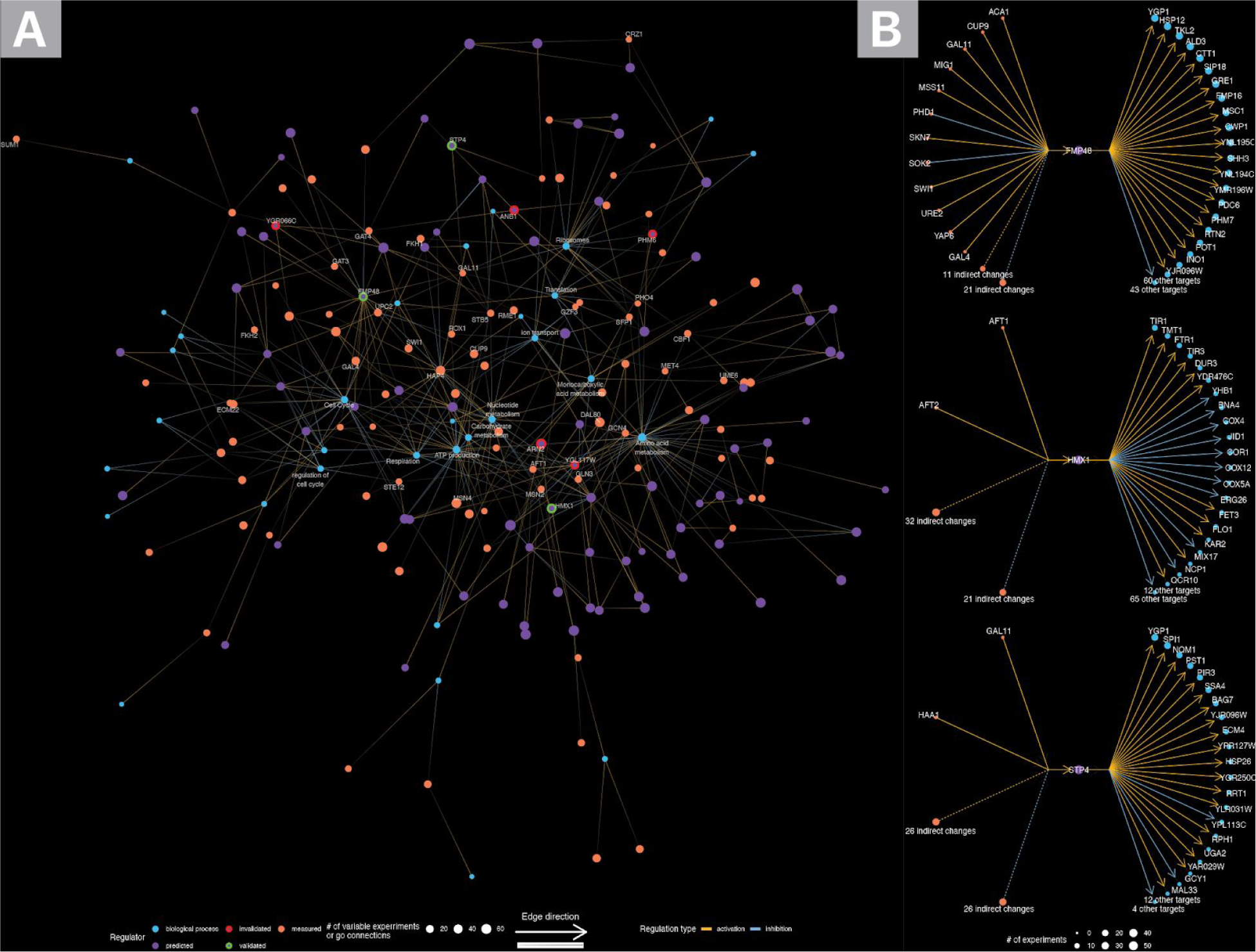
Synthesis of predicted networks. Direct regulation between genes is defined based on causal attribution analysis and indirect regulation of an induced gene is defined if a gene is differentially expressed regardless of whether attribution analysis indicated a direct regulatory relationship. **A.** Edges between both genes with induction experiments and predicted regulators were formed based on regulatory cascades predicted from individual experiments (as shown in Figure 3C). For this visualization, major regulators selected as per Figure 3C are rooted to the induced transcription factor regardless of whether they were directly or indirectly regulated by this gene. Predicted regulators are linked to GO categories based on overlap with their predicted targets and similarly genes with an induction experiment are linked to GO categories based on overlap of either direct or indirect targets with GO categories. **B.** Local networks based on upstream direct/indirect regulators and downstream direct targets of three validated regulators.

### Multiple transcriptional regulators confirmed

Our modeling results highlight many potential new regulators that we sought to confirm experimentally. These regulators include both hubs predicted to regulate targets across many experiments as well as mediators of interesting dynamic phenomena (such as impulses). To validate putative regulators, a separate β-estradiol induction timecourse was generated for each of ten new regulators of interest (Table S4).

Three of these induction timecourses showed strong changes in the putative regulators’ targets (Figure 5, p < 10^-12; overlap of predicted and measured targets by χ^2^ test) (see Supplement). Correctly predicting 3/10 transcriptional regulators is notable, both because few genes are thought to be able to act as specific transcriptional regulators, and because all confirmed regulators were poorly studied. In line with the integrative signals that we aimed to capture in this study (Figure 1E), the three confirmed regulators change in many experiments, and act as hubs which connect diverse TFs to prominent regulatory processes. Each validated regulator temporally preceded its predicted targets and variation in the regulator’s t_rise_ was correlated with changes in its targets’ t_rises_ and the regulator’s v_inter_ was correlated with targets’ v_inters_ (Figure S13). Invalidated regulators, in contrast, either temporally coincided with their spurious targets or were active in a small number of timecourses and thus difficult to distinguish from correlated genes.

**Figure 5.**
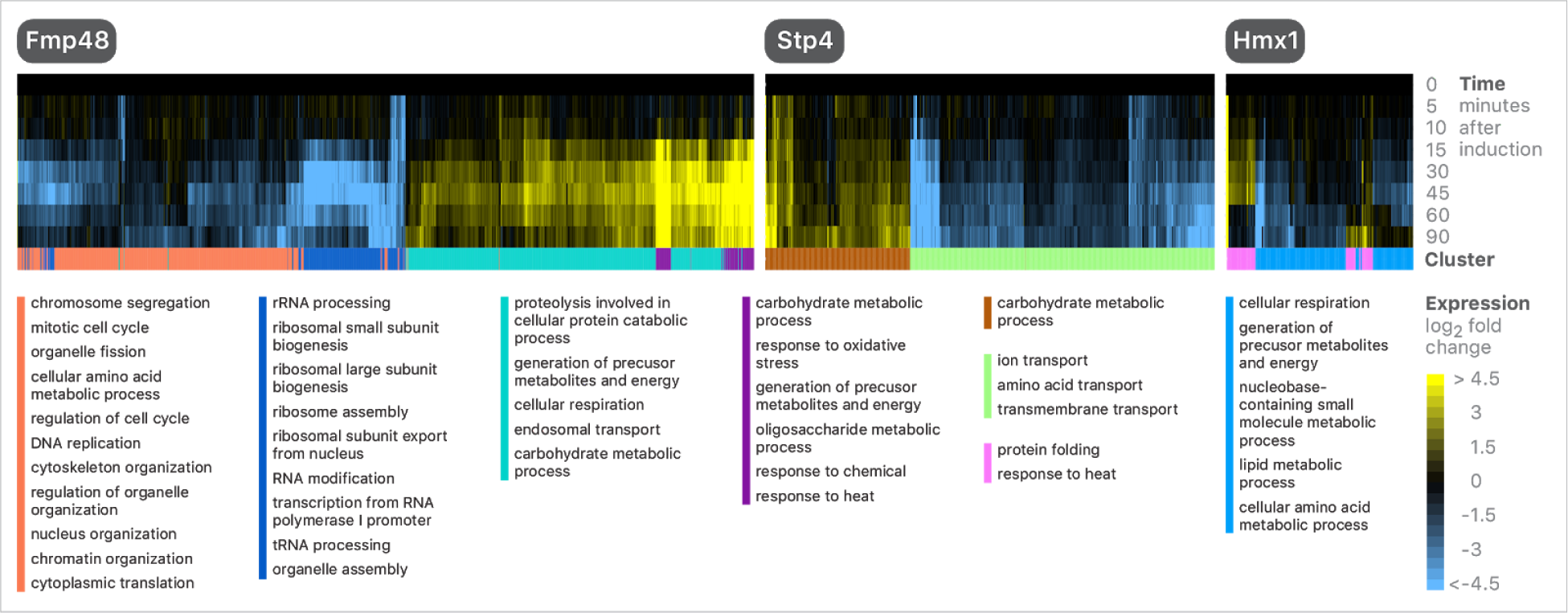
Model-driven identification of transcriptional regulators. All genes passing hard-thresholding in each experiment are shown. K-means clustering was used to cluster responsive genes (K = 4 for Fmp48 and K = 2 for Stp4 and Hmx1) and GO slim gene-sets enriched in each cluster are shown.

Hmx1, notably, mediates indirect effects of Aft1 and Aft2. Induction of Aft1 or Aft2, key regulators involved in iron utilization, results in diverse expression changes. We find that as part of the iron utilization cascade, Aft1/Aft2 activate expression of *HMX1*. Hmx1 induction results in the inhibition of *COX* genes, and genes involved in sterol biosynthesis (a process that requires heme), including *CYB5* (encodes Cytochrome b5) and nearly every *ERG* gene [23], establishing a regulatory link between iron regulation, heme metabolism, and sterol synthesis (Figure 6A). Previously, it was found that *hmx1*∆ cells accumulate heme, suggesting that Hmx1 activity is important role in recycling heme during conditions of iron starvation [24]. *HMX1* induction also represses *NDE1*, which encodes a mitochondrially-localized NADH dehydrogenase and is important for cellular respiration.

**Figure 6.**
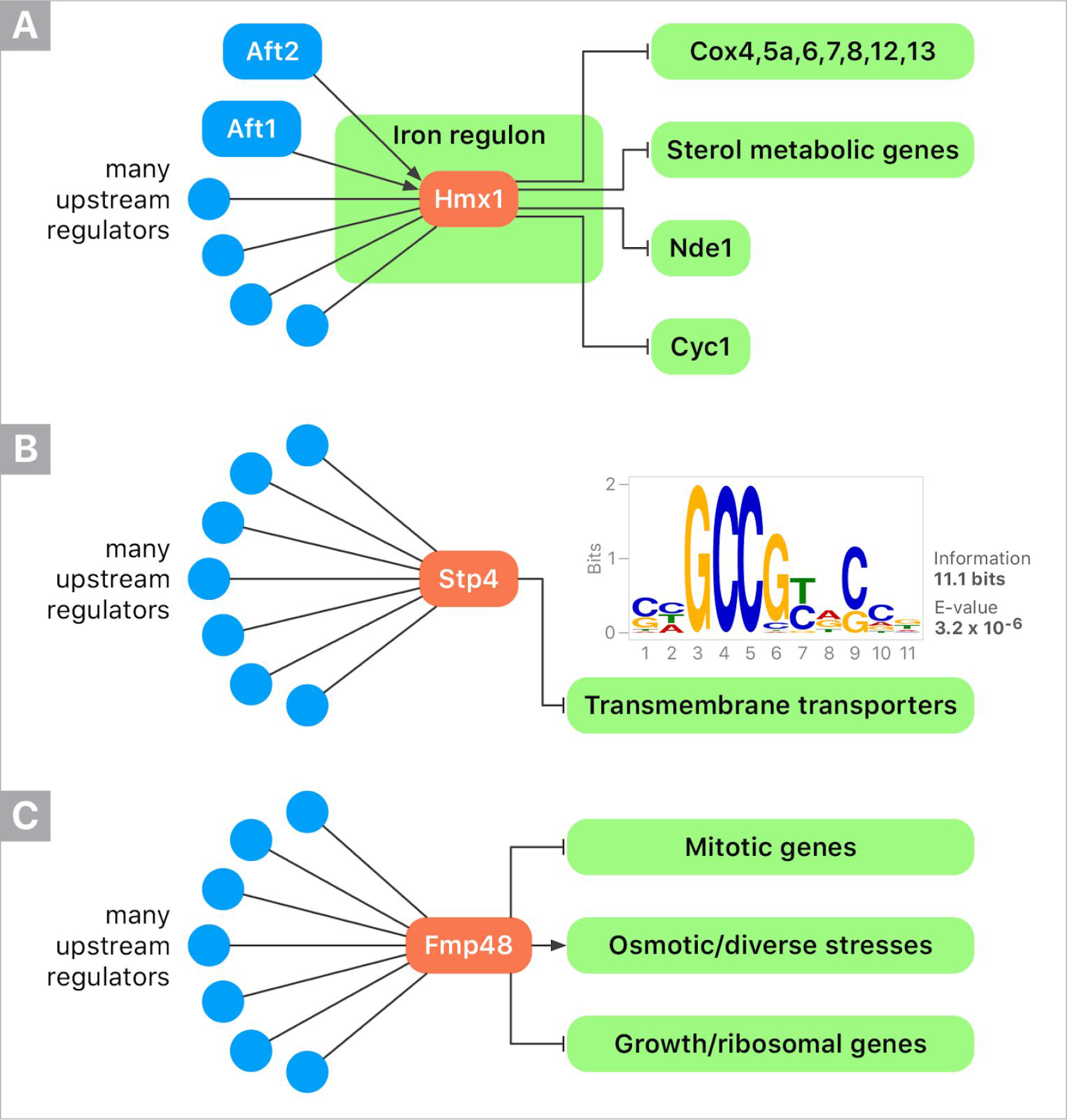
Gene regulatory networks (GRNs) predicted computationally and validated experimentally. TFs with induction experiments used for modeling are shown in blue. Genes that were induced based on model predictions are shown in orange, and responses to those genes are shown in green. **A.** Hmx1 is part of the yeast iron regulon, and upon activation, can repress a set of genes involved in respiration and sterol metabolism. **B.** Stp4 is regulated by a diverse set out TFs. Genes that respond most quickly/strongly to Stp4 induction are repressed, and enriched for amino acid transporters. Promoters of these genes are enriched for the indicated binding motif. **C.** Fmp48 is activated/repressed by many TFs, and in turn, regulates diverse cellular processes.

Stp4 is annotated as a potential TF that contains a Krueppel-like domain [25]. Our data strongly suggest that Stp4 is a *bona fide* TF resulting in activation/repression of hundreds of genes (Figure 5). In the Stp4-responsive gene set there is a set of 83 fast-responding, strongly repressed genes. Using MEME, we determined that promoters of these genes are enriched for the motif GNRCGGCY; this motif is nearly identical to a Stp4 motif derived from a protein binding microarray [26]. Genes responsive to this motif are strongly enriched for transmembrane transport (corrected p-value = 2.07 x 10^−8^), and include the biotin transporter (*VHT1*) and heme transporter (*PUG1*) (Figure 6B). This cluster contains genes strongly inhibited by Stp4, including numerous amino acid transporters (*GAP1, TAT1, PUT4, ODC1, BAP3, YCT1,* and *GNP1*).

Fmp48, named after “found in mitochondrial proteome”, is a putative protein of unknown function with predicted kinase activity. In the Fmp48 validation experiment, ~1800 genes changed by more than two-fold, making Fmp48 a prominent transcriptional regulator. Genes that respond to Fmp48 activation fall into three clear classes: activation with fast kinetics, repression with fast kinetics, and repression with slow kinetics (Figure 5). Quickly repressed genes are mostly strong enriched for genes involved in rRNA processing, while slowly repressed genes are strongly enriched for genes involved in mitotic cell cycle and DNA replication (corrected p-values < 0.001); in fact, the class of slowly repressed genes includes all of the core histones: H3 (*HHT1* and *HHF1*), H4 (*HHT2* and *HHF2*), H2A and H2B (*HTA1*, *HTA2*, *HTB1*, and *HTB2*), as well as H2A-Z *(HTZ1*), Cen H3 (*CSE4*), and H1 (*HHO1*). Comparing our data to a previous gene expression profiling study of ~1400 deletion mutants [27], we find that the Fmp48 45-minute timepoint is most correlated with *cst6*Δ, *ram1*Δ, and csm1Δ (with Pearson correlation of ~0.2) and most anti-correlated with *nrm1*Δ, *hur1*Δ, and *ecm30*Δ (with Pearson correlations of ~ −0.15). *NRM1* encodes a transcriptional co-repressor of cell cycle genes, and indeed, we find that Nrm1 strongly represses cell cycle genes when induced (Figure S14). This suggests that Fmp48, as part of its regulon, may activate Nrm1 activity to inhibit cell-cycle progression.

Fmp48 is a regulator of metabolic and diverse stress-responsive genes (Figure 6C). Previously, it was found that Fmp48 interacts with TOR, and that overexpression results in alterations in mitochondrial morphology and differential growth phenotypes depending on carbon source (slight overexpression of Fmp48 is toxic when glycerol is a carbon source, but not when raffinose is) [28]. Genes that are activated by >2-fold in both our dataset and a previously published Fmp48 overexpression gene expression dataset [28] are most enriched for methylglyoxal metabolic processes (corrected p-value 1.12 x 10^−7^) and cellular response to chemical stimuli (corrected p-value 2.39 x 10^−7^) (Figure S15). The most up-regulated genes across both datasets includes *HSP26, SIP18*, *ALD3*, *FMP16*, *CTT1*, *GND2 NQM1*, *GRE1*, *HSP12*, *SPI1*, *SSH3*, *DDR2*, *SOL4*, *RTC3*, *MSC1*, *RTN2*, *PAI3*. *GRE1* and *SIP18* are paralogs that are important for overcoming the dehydration-rehydration process, and *CTT1* encodes catalase, a potent antioxidant. Beyond metabolic and stress-responsive genes, this gene set includes *RTN2*, which encodes a gene important for maintaining Endoplasmic Reticulum shape and has stress-dependent localization [29]. Clearly, our data highlight Fmp48 as a prominent, hub-like regulator.

## Discussion

In order to understand regulatory architecture, we require datasets that elicit diverse physiological regulatory responses, and possess sufficient information to disambiguate the drivers of each regulatory response. Synthetic biology holds great promise for creating such datasets, and, when combined with new analytical tools, can be utilized to identify new regulators and GRNs.

Here, we demonstrated the power of using large datasets and TF induction timecourses to reveal new regulatory connections. Four results are worth highlighting. First, our dataset reveals homeostatic relationships. By inducing a TF and measuring changes in expression over time, direct activation/repression can be revealed, and novel cases of feedback can be observed as impulse-like transcriptional responses. Second, expression variation is associated with a modest number of transcriptional regulators with major effects, while many perturbations elicit minimal transcriptional responses; in fact, ~40 of the TF induction experiments (~20% of all experiments) elicited no significant changes in gene expression (Figure S12). Third, using kinetic information is essential for prioritizing potential causal regulatory relationships. By integrating hundreds of time courses with a dynamical systems model that explicitly includes time, we can make predictions of new regulatory interactions without the use of prior knowledge. Fourth, we highlight the regulatory potential of previously under-appreciated regulators of gene expression.

Approaching genome-scale modeling of network regulation from dynamic data also yielded insights about how to collect, process, and analyze such data. Because most genes do not respond in a typical induction experiment, we used hard-thresholding to remove (i.e., set equal to zero) the majority of values in our dataset, leaving ~100,000 timecourses with coherent, biologically-feasible patterns of variability. Having first identified signal-containing timecourses, we were then able to determine the nature of this signal using parametric models and by modeling the dataset-level relationships between signals. When fitting a genome-scale regression model we made a number of assumptions about biological processes to include versus those we should ignore based on our experimental design. In doing so, our model may fail to capture a number of important regulatory phenomena including complex combinatorial regulation, post-transcriptional regulation, post-translational regulation, localization and regulation due to non-proteins (e.g., metabolites) [30–34]. These phenomena could hamper our analysis to the extent that regulators are absent, their concentrations are misrepresented, or their kinetics are temporally shifted. Because a transcriptional model is inherently incomplete, our modeling approach was structured to be robust to mis-specification by describing variables directly from data rather than creating latent variables. Our model builds relationships between genes with coherent regulatory relationships without being grossly biased by regulation that it cannot represent. This model is inherently incomplete, and accordingly, our model only explains ~40% of expression variation (Figure S11). An additional challenge we faced when fitting a model that allows for regulation by any gene is distilling a single regulator from a set of possibly highly-correlated possibilities. Some regulators not well-represented by our dataset may also be correlated with measured transcripts raising the possibility that predicted transcriptional regulators may be false-positives if they correlate to an unmeasured regulator. By utilizing >200 timecourses, we are in a regime where identifiability of an individual regulator becomes possible, because we are able to break the correlation between pairs of genes (Figure S10). It is also noteworthy that this ability decays quickly once an appreciable fraction of timecourses are removed (Figure S10).

In a given timecourse, genes with similar kinetics often have a common regulator, and we expect kinetic similarity between genes to be reflected in the DNA composition of their promoters. More could be done here, perhaps by training a model with grouped regularization [35] based on either existing motifs (see [18]) or motifs directly learned from the data [36]. In practice, some transcriptional regulators (e.g., chromatin modifiers) will not affect targets in a manner consistent with an easily discoverable DNA motif, and in other cases global effects may cause widespread expression changes [37]. Motifs, nevertheless, may kinetically localize despite the absence of clear direct regulation by the induced TF. For example, ribosomes that are acutely inhibited in the Aft1 and Pho4 experiments are enriched for the PAC sequence motif but there is little expression evidence that the associated regulators are operating. To avoid the case where we are over-fitting to known relationships, we opted to build an *ab initio* model of transcription which allowed for regulatory relationships between any pairs of genes and fit based on coherent patterns of expression. Only one of the three validated transcriptional regulators was annotated as a putative TF. For the remaining non-TFs, other downstream regulators may ultimately drive variation (whether by changes in a secondary messenger, TF localization, etc.). Still, in each case the predicted regulator is sufficient to elicit a regulatory response and is more proximal to the effect than the induced TF, and thus more informative of direct regulation.

In the future, improvements in synthetic biology and computational modeling will result in even better predictive models. In the approach presented here, only one TF was perturbed at a time, resulting in a large but relatively sparse gene expression dataset. The use of combinatorial perturbations, as well as induction of non-TFs, will result in richer dynamic datasets. As new timecourse datasets become available and are integrated with time series analysis and prior knowledge, we expect predictive models to require fewer experiments to build. Moreover, dataset generation and model evaluation naturally dovetail when using synthetic perturbations. Regulators can be easily tested with new induction experiments. When modeling predictions succeed, we confirm new biology; when they fail, the model gets better.

## Acknowledgments

We thank Michelle Dimon, Ali Bashir, Chiraj Dalal, David Botstein, Dan Gottschling, Rochelle Buffenstein, and John Platt for comments and critical feedback on the manuscript.

---

**Figure S1:**
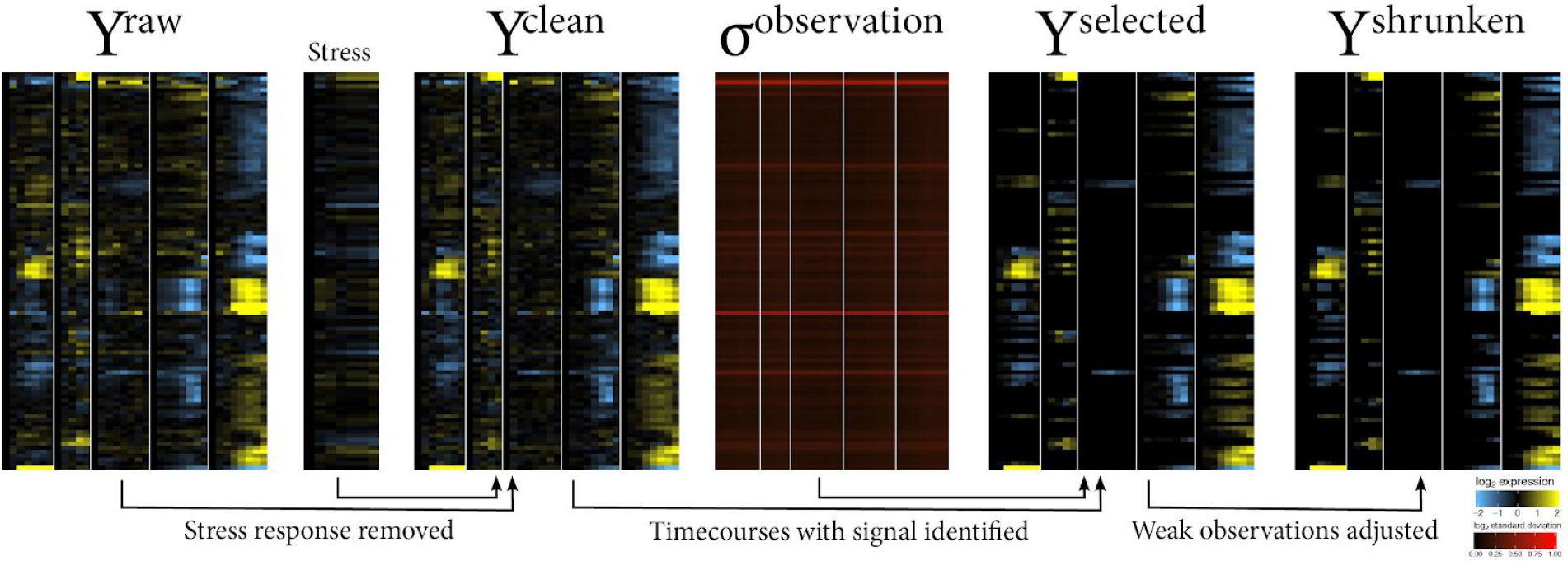
Systematic identification and summarization of regulatory signals. Since most transcriptional regulators affect a relatively small number of target genes, meaningful changes in expression are relatively sparse (~9% of timecourses). These signal-containing timecourses are distinguished from timecourses which are purely noise, by first regressing out an average stress response, then selecting timecourses with extreme observation-level signal-to-noise and finally shrinking observations towards zero based on signal-to-noise.

**Figure S2:**
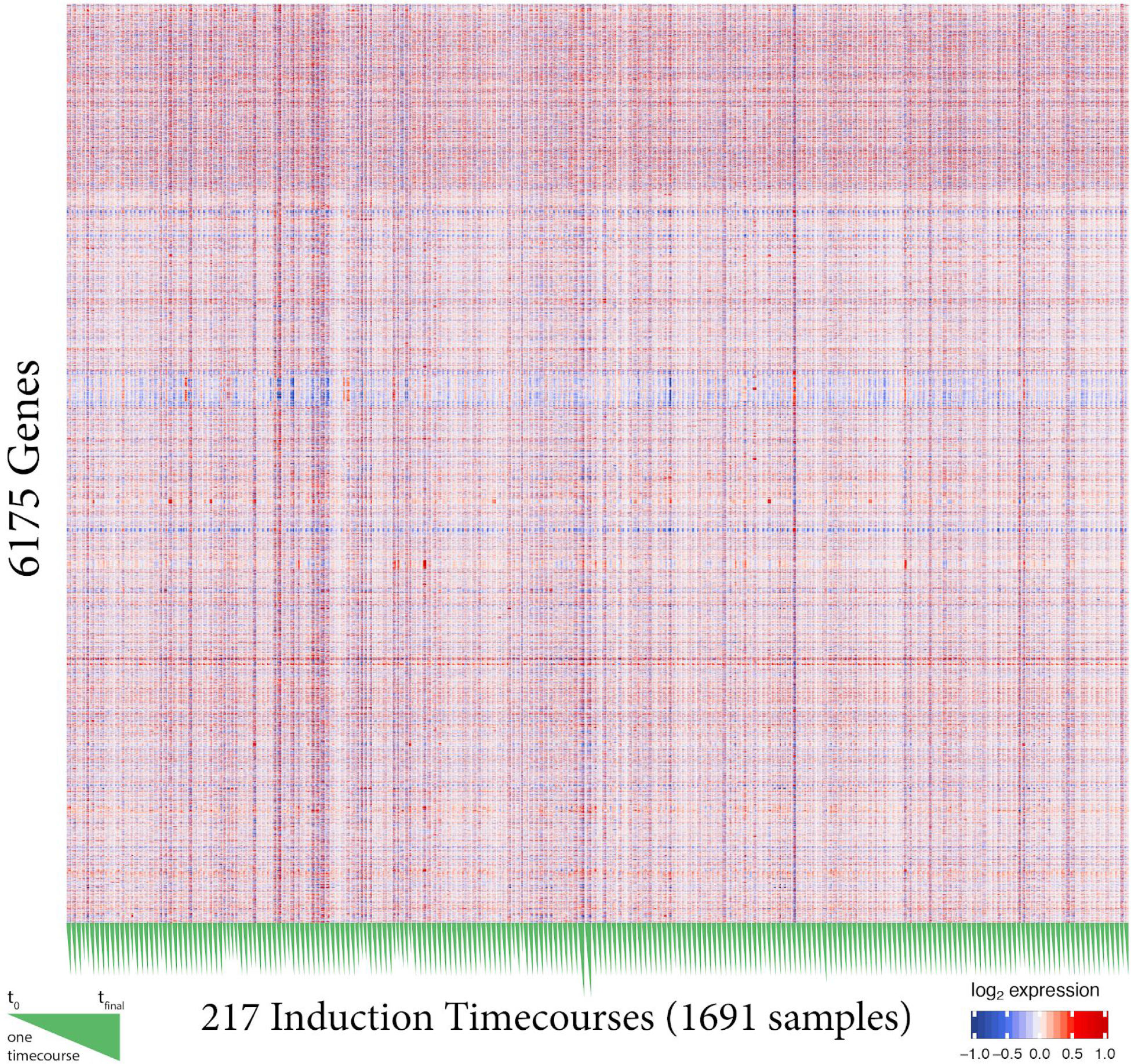
Full transcriptome of the “raw” gene expression data. Genes are sorted alphabetically from bottom to top. Note the increase in variability of gene expression for genes near the top of the clustergram (these are mostly genes that begin with “Y” as they have no known function and lack a standard name).

**Figure S3:**
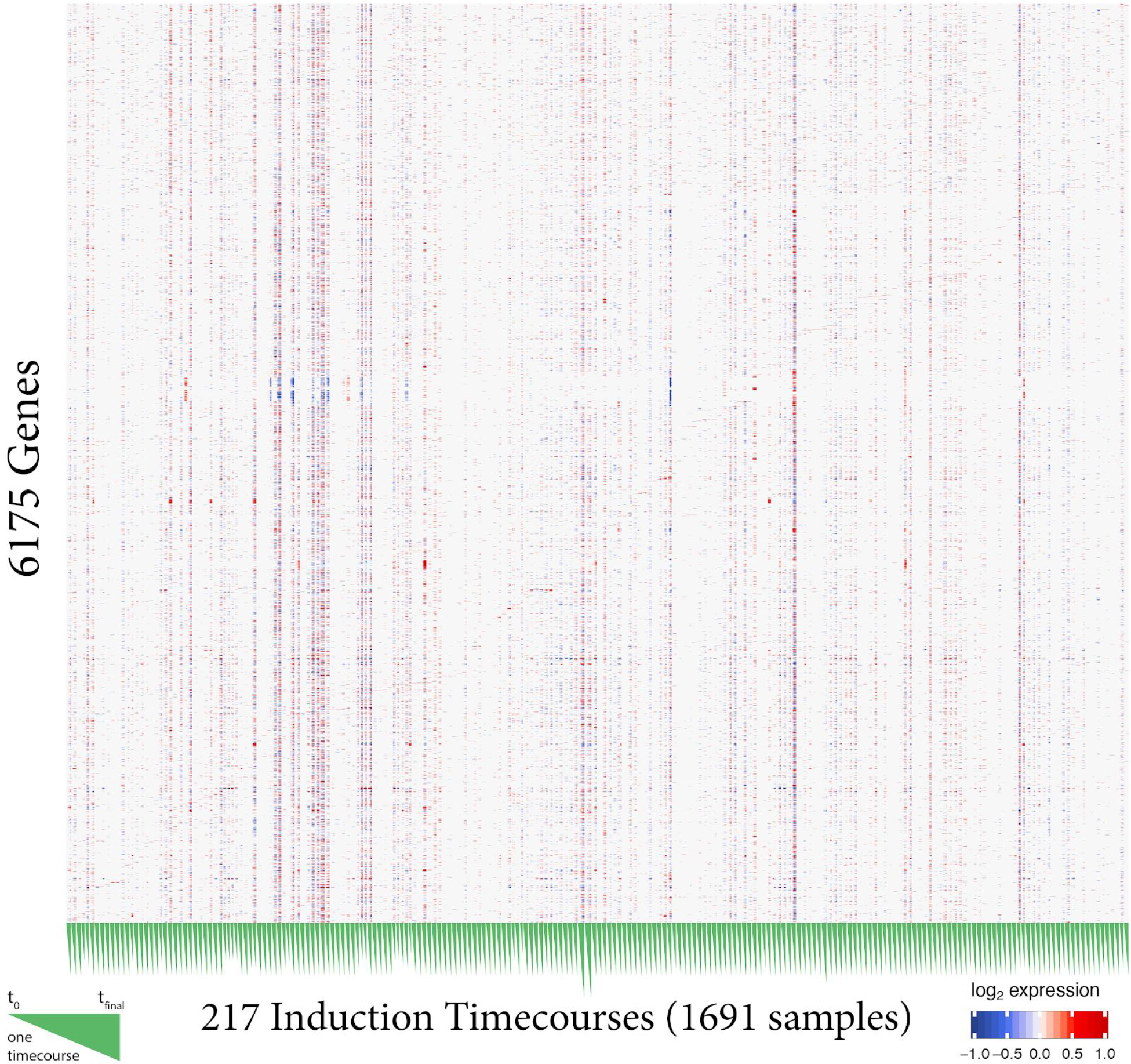
Full transcriptome of the “shrunken” gene expression data. Genes are sorted alphabetically from bottom to top.

**Figure S4:**
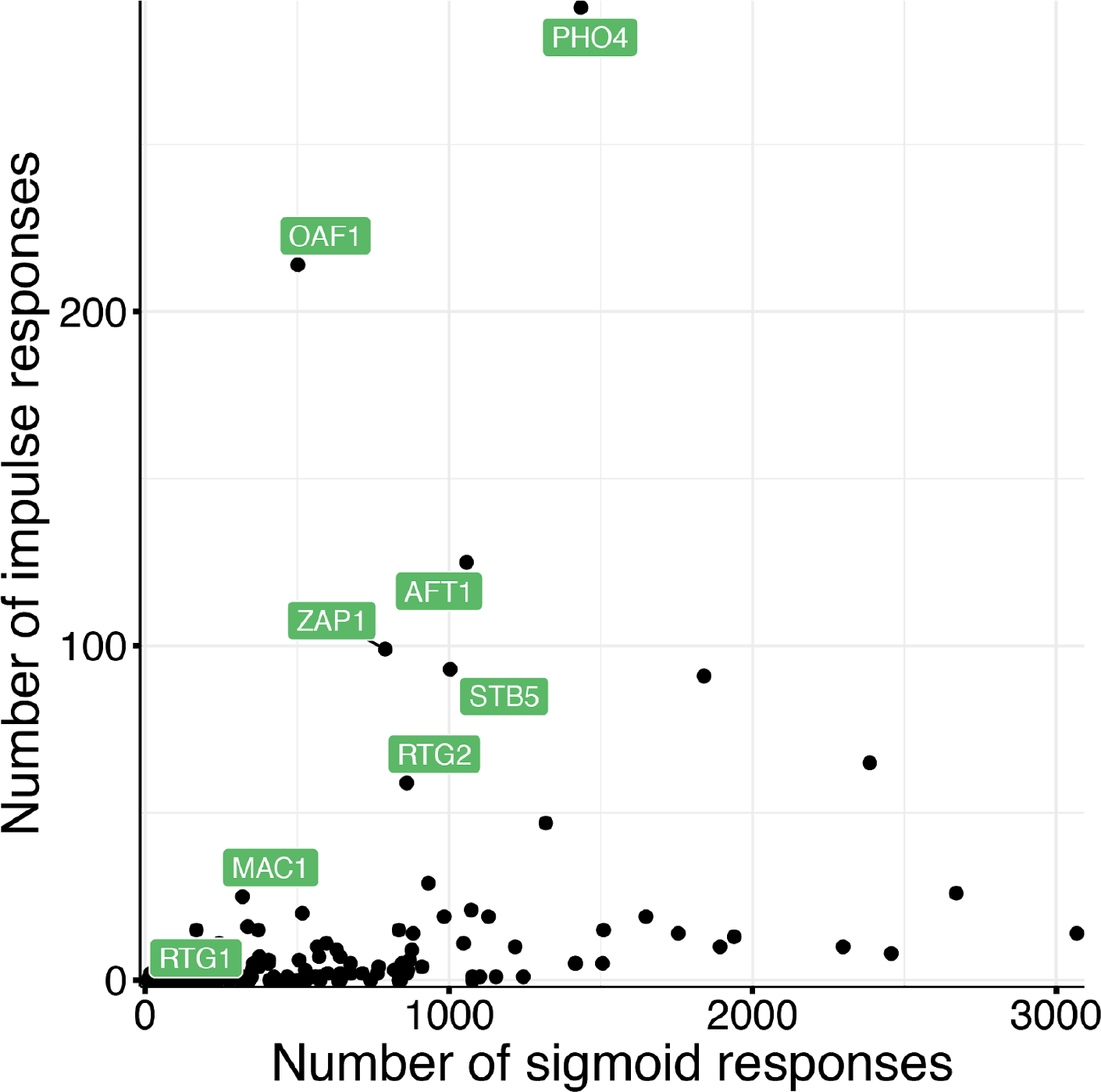
Counts of impulse vs. sigmoids across experiments. Transcriptional regulators are characterized based on the number of transcriptional responses that are sigmoid (e.g., turn on) versus impulses (e.g., turn on, then off). Pho4, Oaf1, Aft1, and Zap1, among other transcription factors, are highly enriched for impulse dynamics.

**Figure S5:**
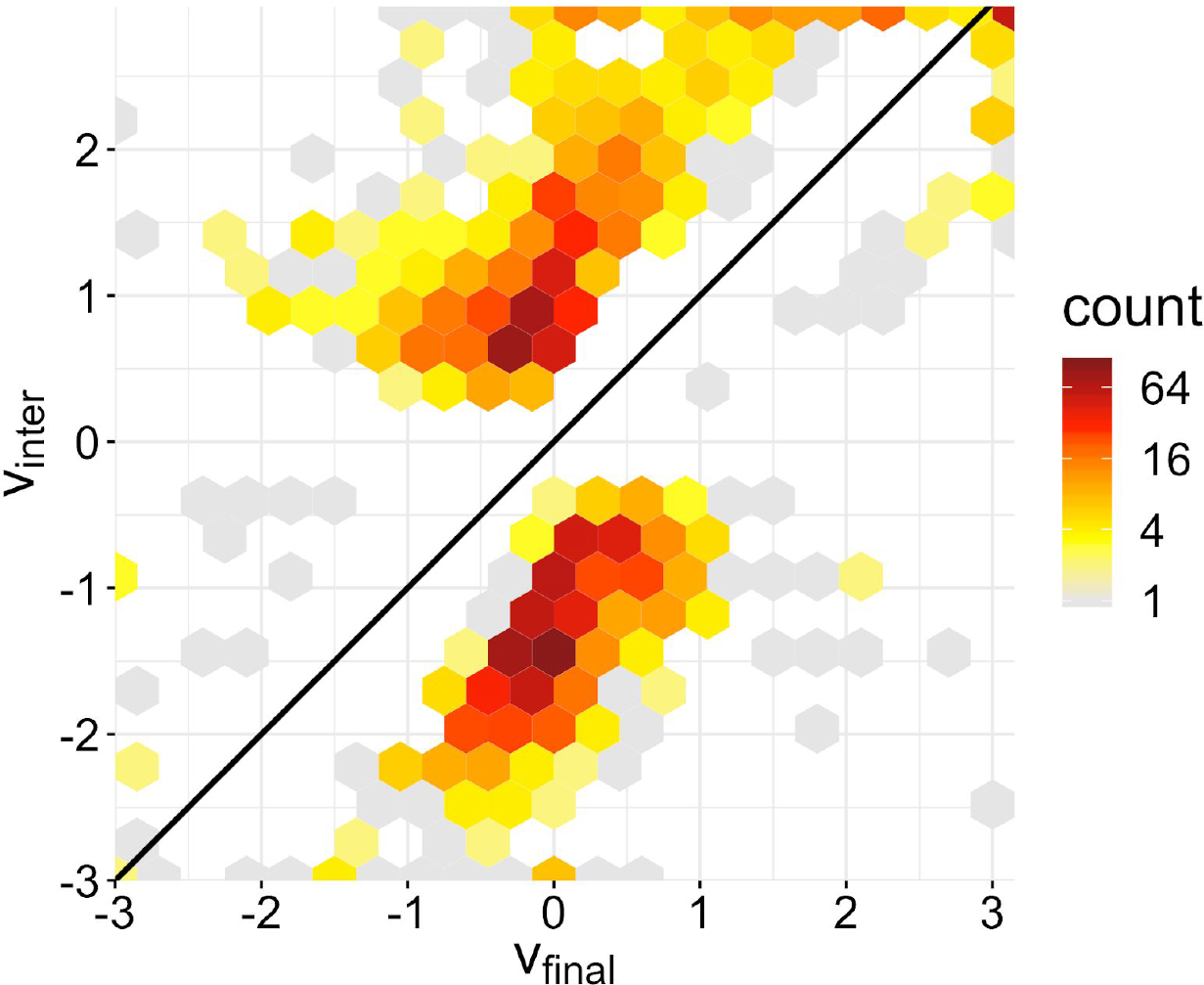
Most impulses exhibit near-perfect adaptation. v_inter_ is compared to v_final_ for all timecourses exhibiting impulse dynamics. The absolute value of each coefficient was floored to three for visualization.

**Figure S6.**
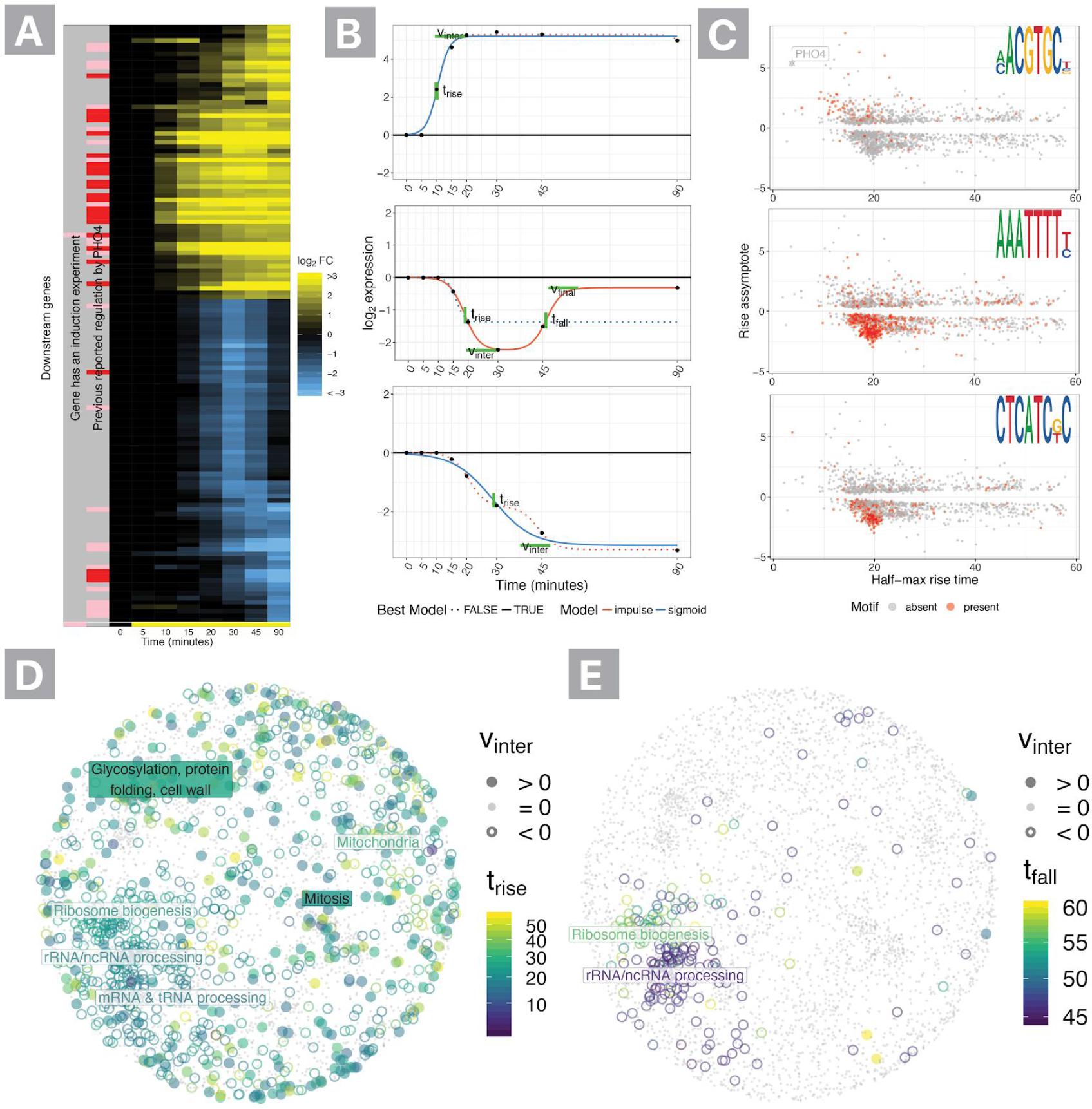
Diverse, functionally significant regulation in the Pho4 induction experiment. **A.** Heatmap summary of Pho4 induction timecourse showing all >4-fold changes. **B.** Parametric summaries of representative sigmoidal activation and inhibition timecourses and impulse (double sigmoid) modeling of transitory inhibition. Sigmoids are summarized by half-max time (t_rise_) and the asymptote (v_inter_), while impulses include a second half-max (t_fall_) time and final assymptote (v_final_). The strongest supported model for each timecourse is shown as a filled in line, while the alternative model is shown with a dashed line. **C.** K-mers enriched in the promoters of regulated genes are overlaid on summary of each gene’s t_rise_ and v_inter_. **D.** Response kinetics are overlaid on gene coordinates based on synthetic lethality as a surrogate for functional similarity. Pho4 rapidly inhibits ribosome biogenesis, rRNA processing and mRNA processing. **E.** The rRNA processing and ribosome biogenesis responses are each acute inhibition impulses that can be clearly distinguishing based on kinetics.

**Figure S7:**
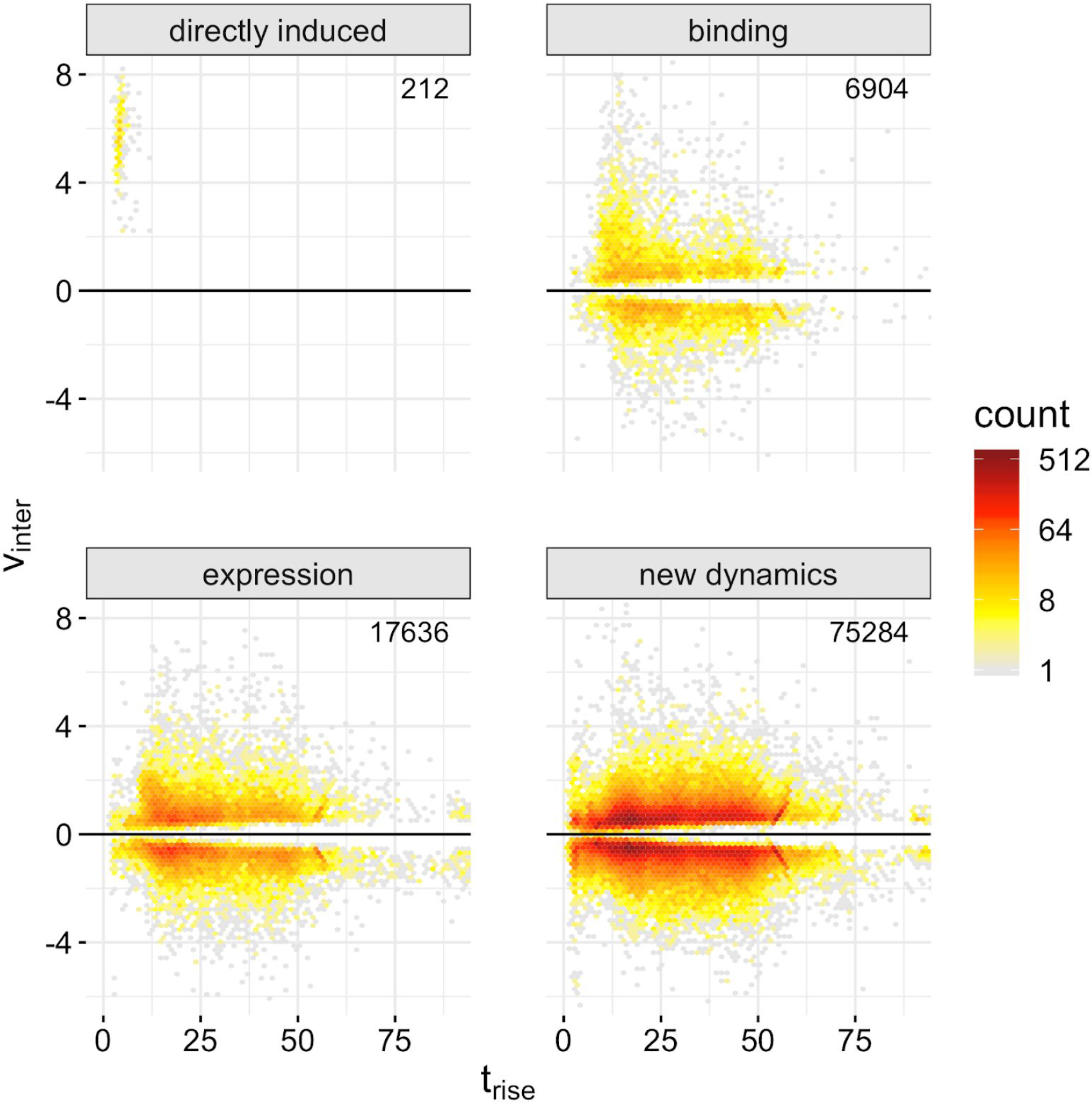
Functional classes of kinetic responses. For each of the >100,000 timecourses with parametric fits, timecourses were divided into four categories: directly induced (TF induced in cognate experiment), binding (direct regulation based on direct-binding data from Yeastract), expression (co-expression of TF and responsive gene based on data from Yeastract), and new dynamics (newly discovered gene associations outside the other three classes).

**Figure S8:**
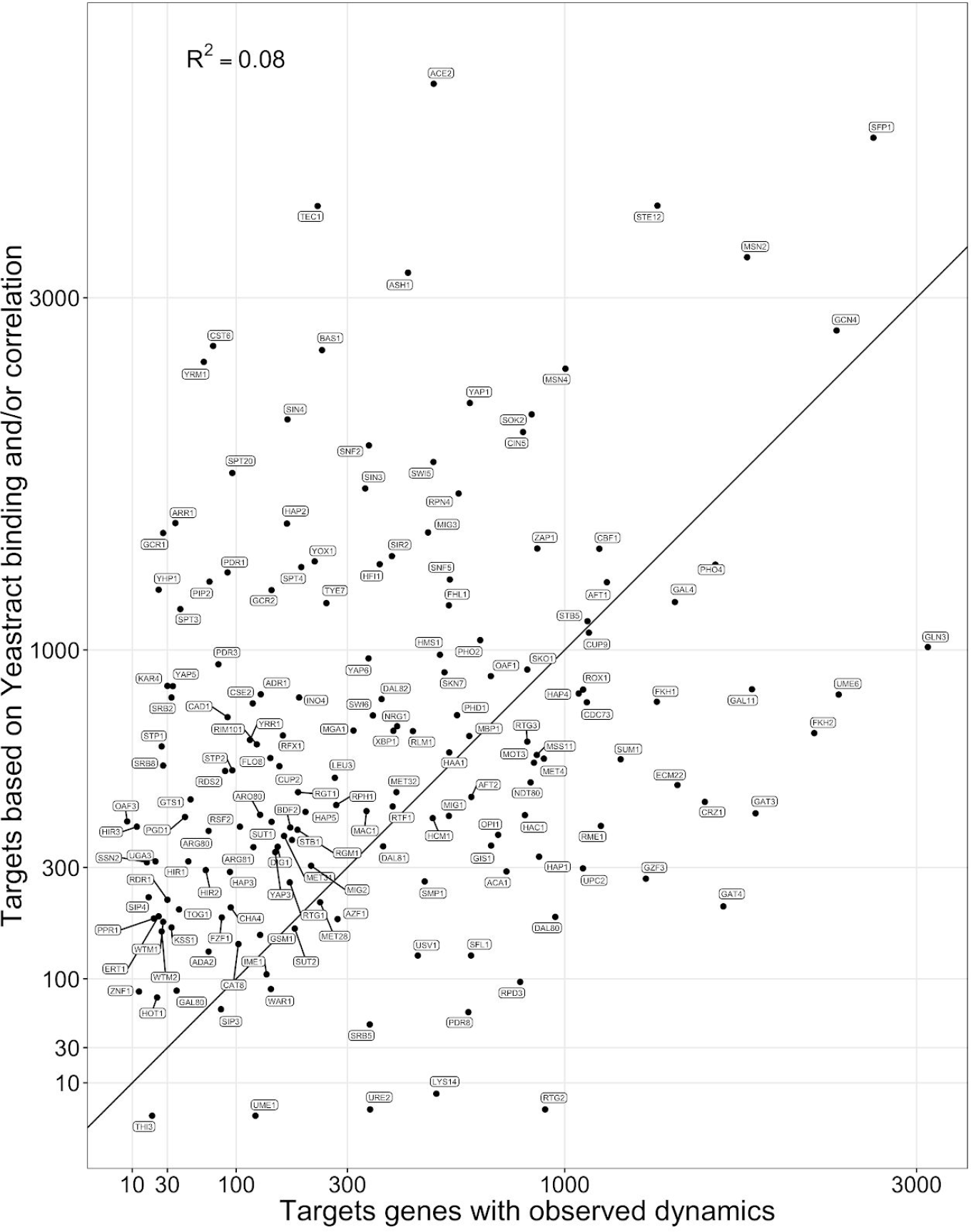
Response magnitude comparison with reported regulation. Scatter plot of number of targets or correlation-associated genes (Yeastract) versus the number of genes with significant dynamical responses in the “shrunken” data.

**Figure S9:**
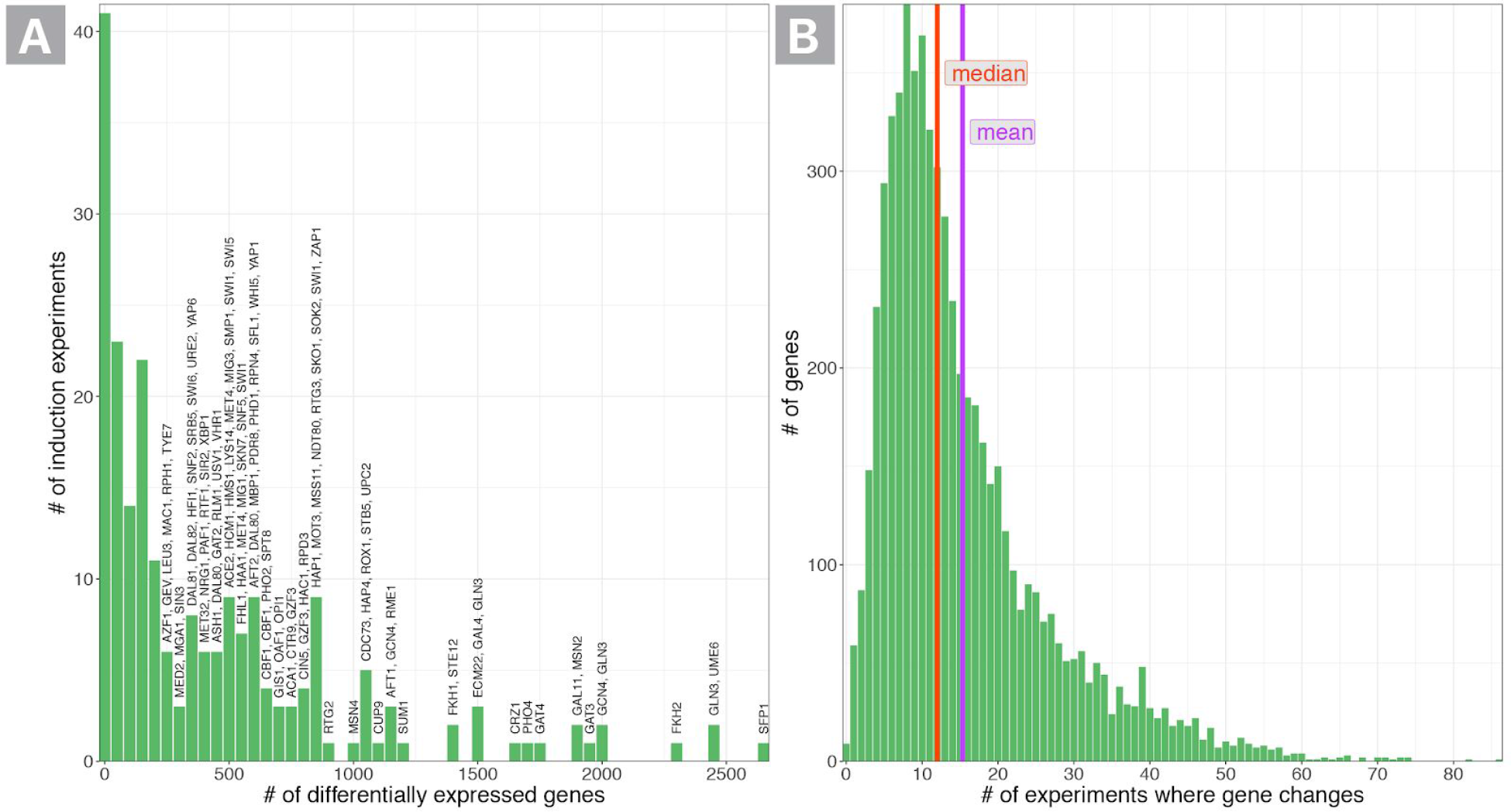
Extent of differential expression per experiment or gene. A) Histogram of the number of differentially expressed genes in each experiment. B) Histogram of the number of transcriptional regulators under which a gene changes (median 12, mean = 15.3, maximum would be changing under all 203 distinct induction experiments).

**Figure S10:**
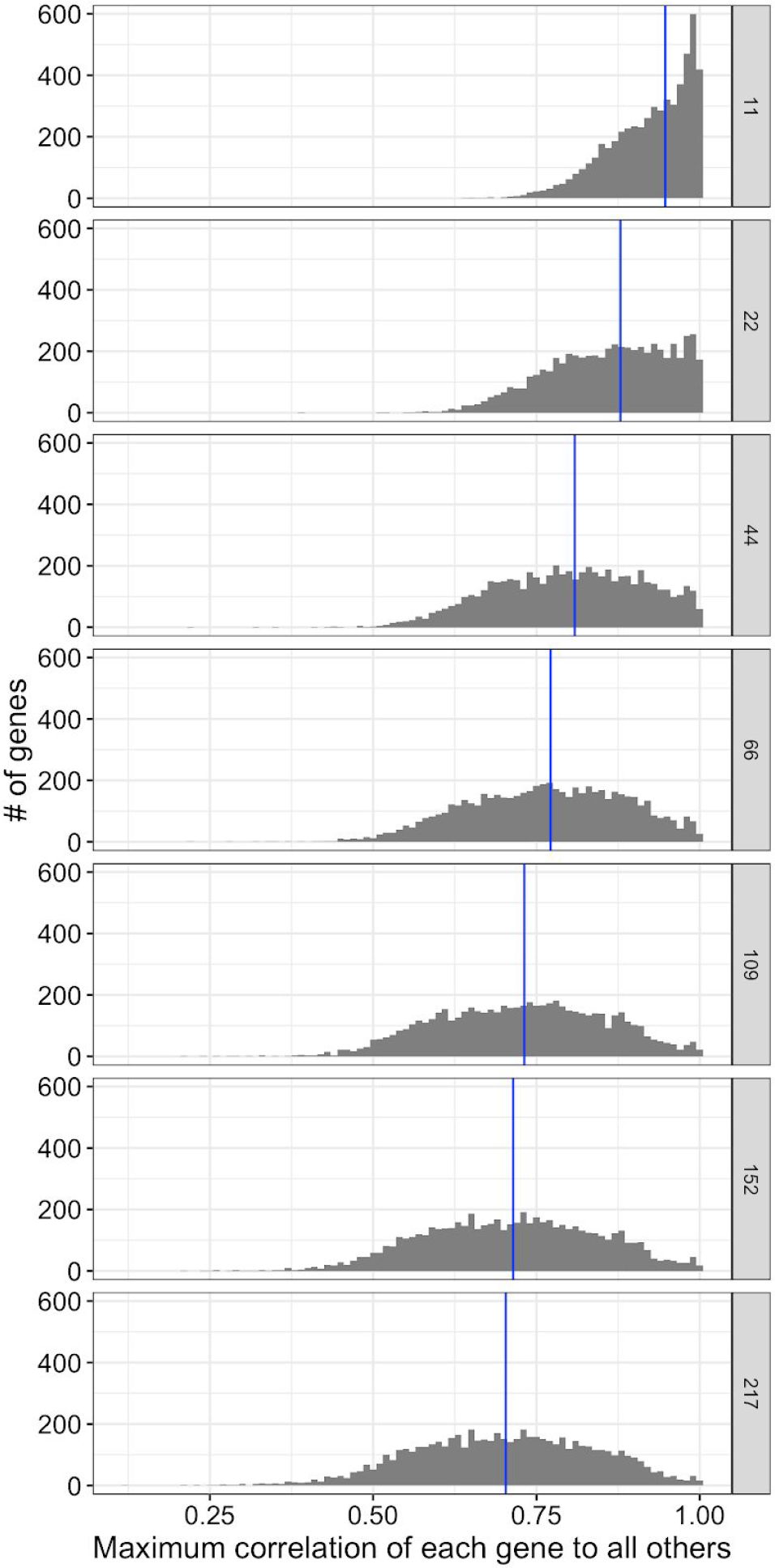
Histograms of maximum correlation of genes to all other measured genes are shown as experiments are pruned from the dataset. Panels indicate the number of experiments included in the analysis, where smaller datasets retain the experiments with the largest number of differentially expressed genes. Each gene is summarized based on the maximum correlation of its expression across all included experiments and timepoints to every other genes’ expression. The blue line indicates the median of the maximum correlation of genes. For the purpose of calculating medians, genes which are dropped when constructing the reduced datasets are represented with a correlation of one (since they are impossible to discriminate from other absent regulators).

**Figure S11:**
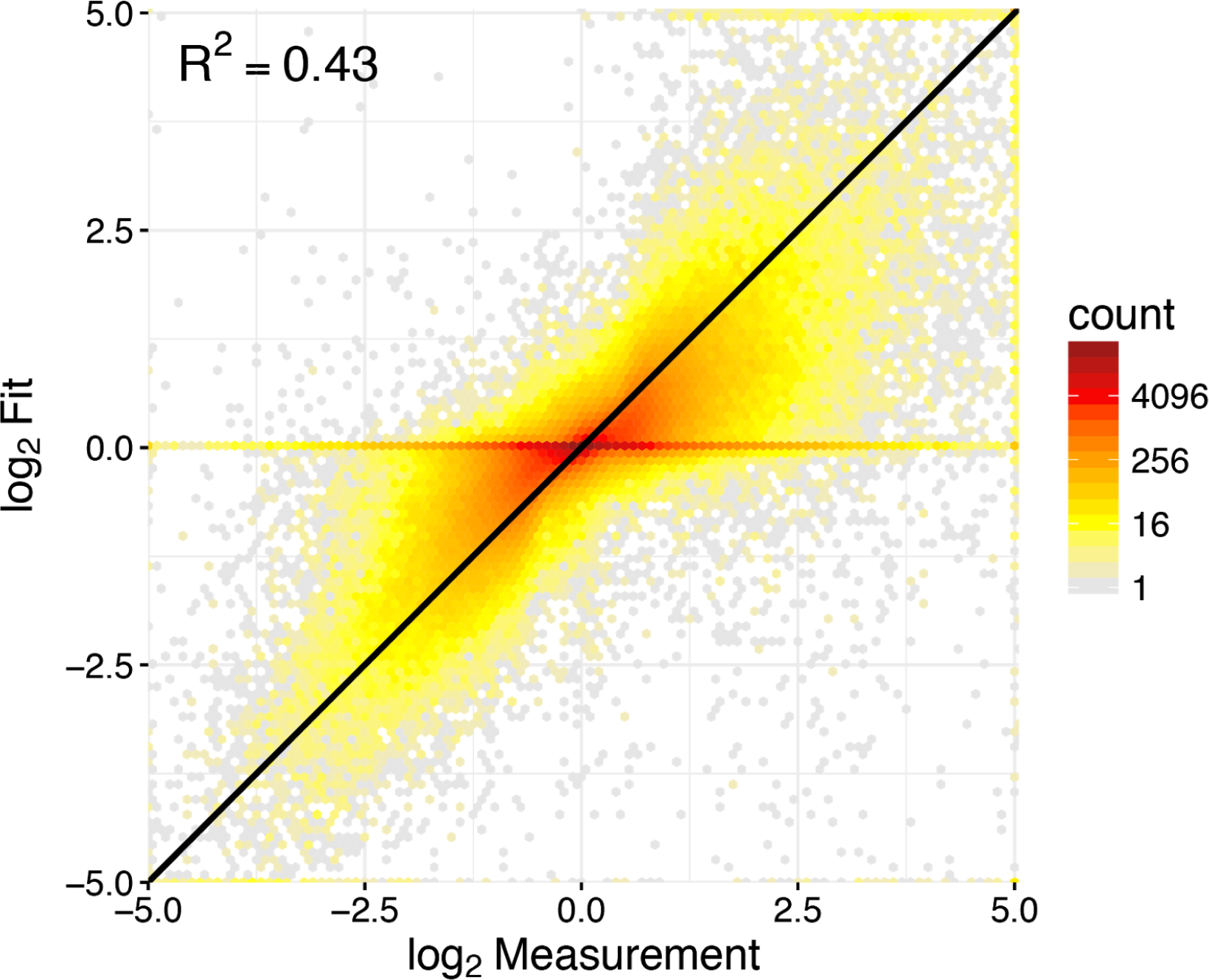
Fit of regulatory model to observed gene-expression measurements. Each observation compares measured log2 fold-changes from the dataset that the whole-cell model was fit to (i.e., time courses that passed full noise model and filters; see section 8 of the supplement) with the fitted fold-changes predicted from the whole-cell model.

**Figure S12:**
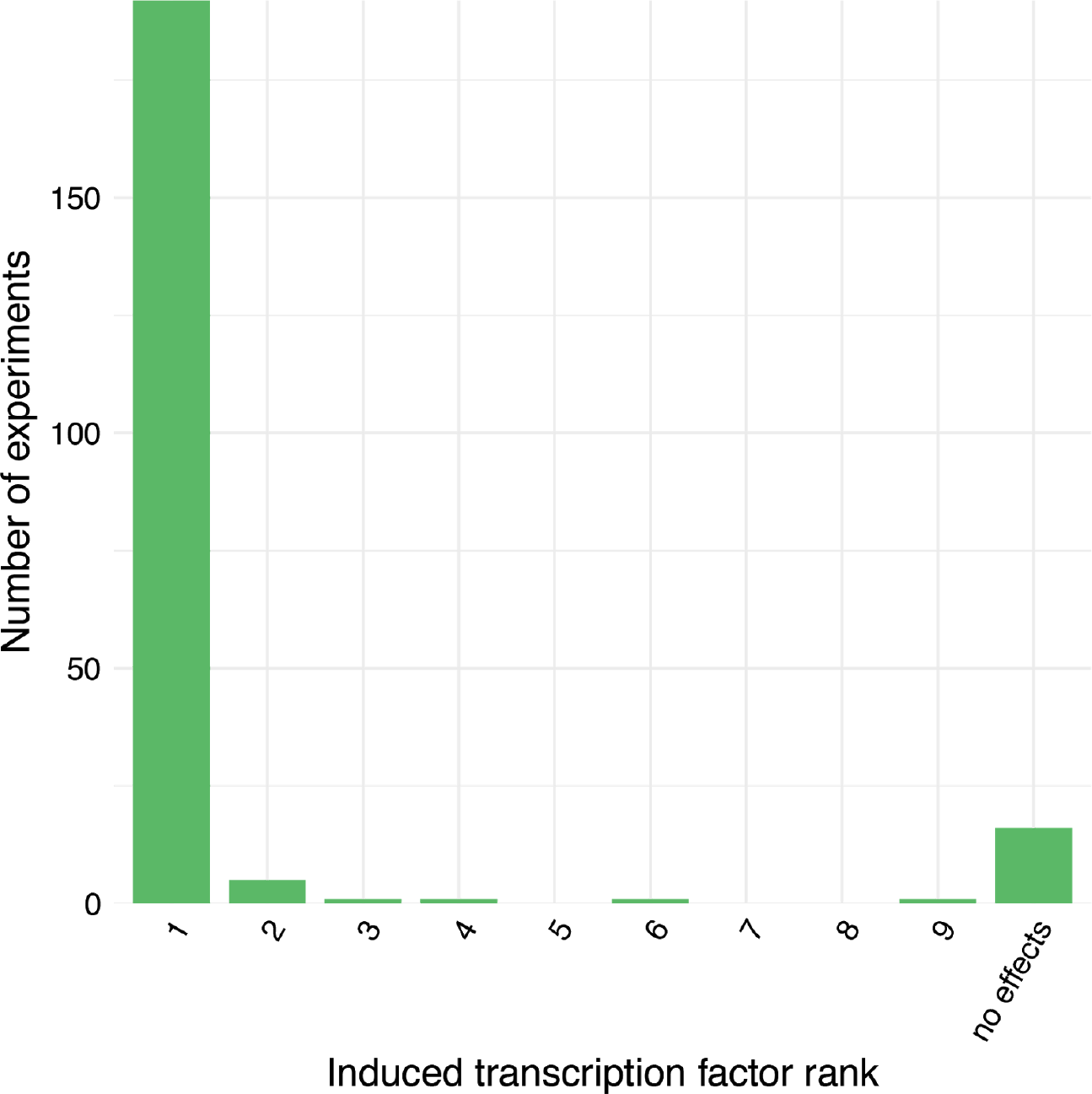
Induced transcriptional regulators are the primary drivers of gene expression changes in most experiments. Each experiment summarized differentially expressed genes based on the regulator with the largest attributed role in achieving the rise time. The rank of the induced transcription factor among regulators, derived from the model, is shown across all experiments.

**Figure S13:**
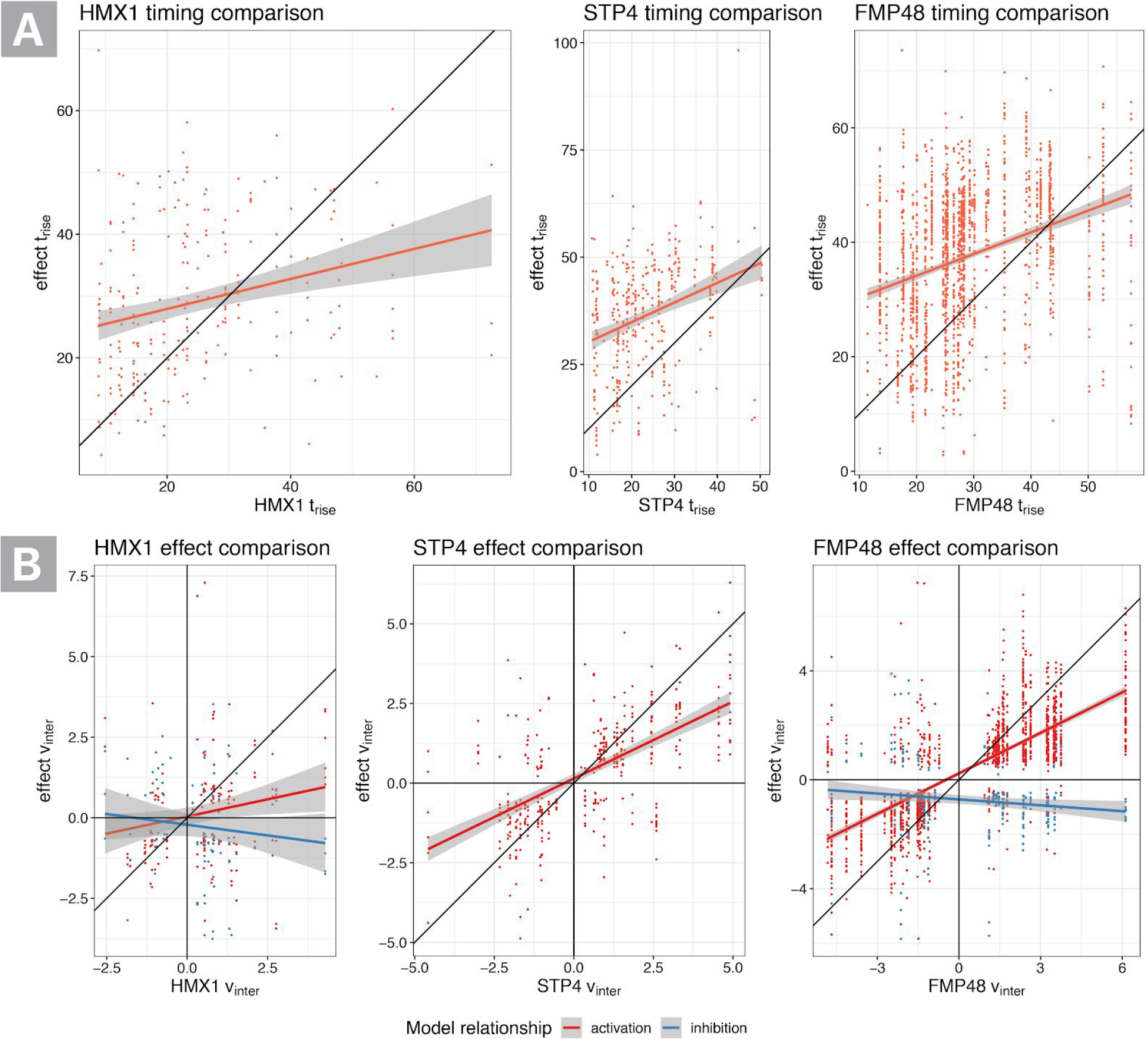
Identifying latent transcriptional regulatory hubs. **A.** For each gene whose expression change is partially attributed (attributing at least 5% of variation) to a regulator’s levels in 5 or more experiments, the timing of the regulator is compared to its predicted targets within the same experiment. **B.** The v_inter_ value (i.e., expression-level asymptote) of each regulator is compared to the v_inter_ of each of its effects in the same experiment. Targets are colored based on whether the regression model indicates an activating or inhibitory relationship.

**Figure S14:**
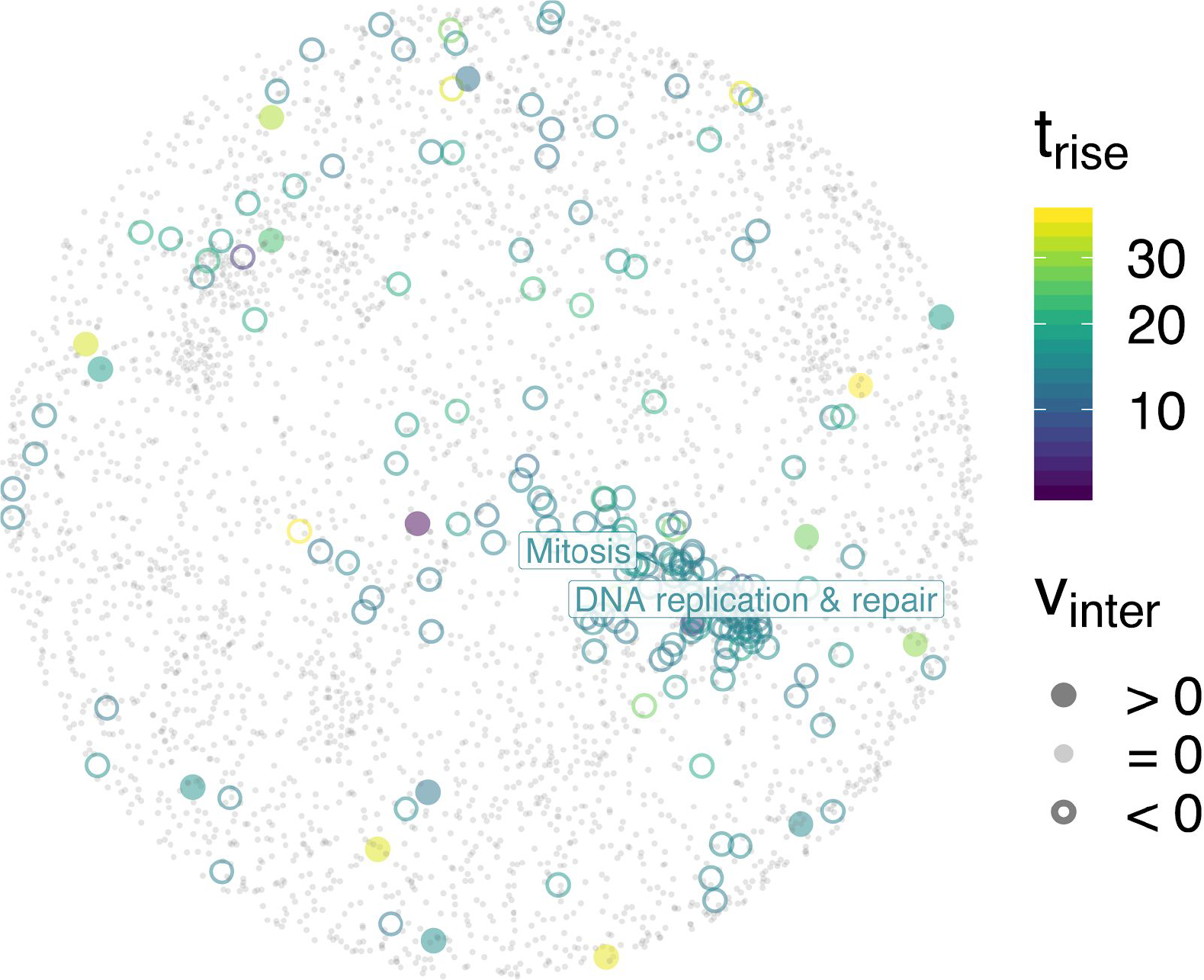
Transcriptional response to *NRM1* induction mapped onto the genetic interaction network. Genes kinetics are summarized based on timing and direction of change as per Figure 2D.

**Figure S15:**
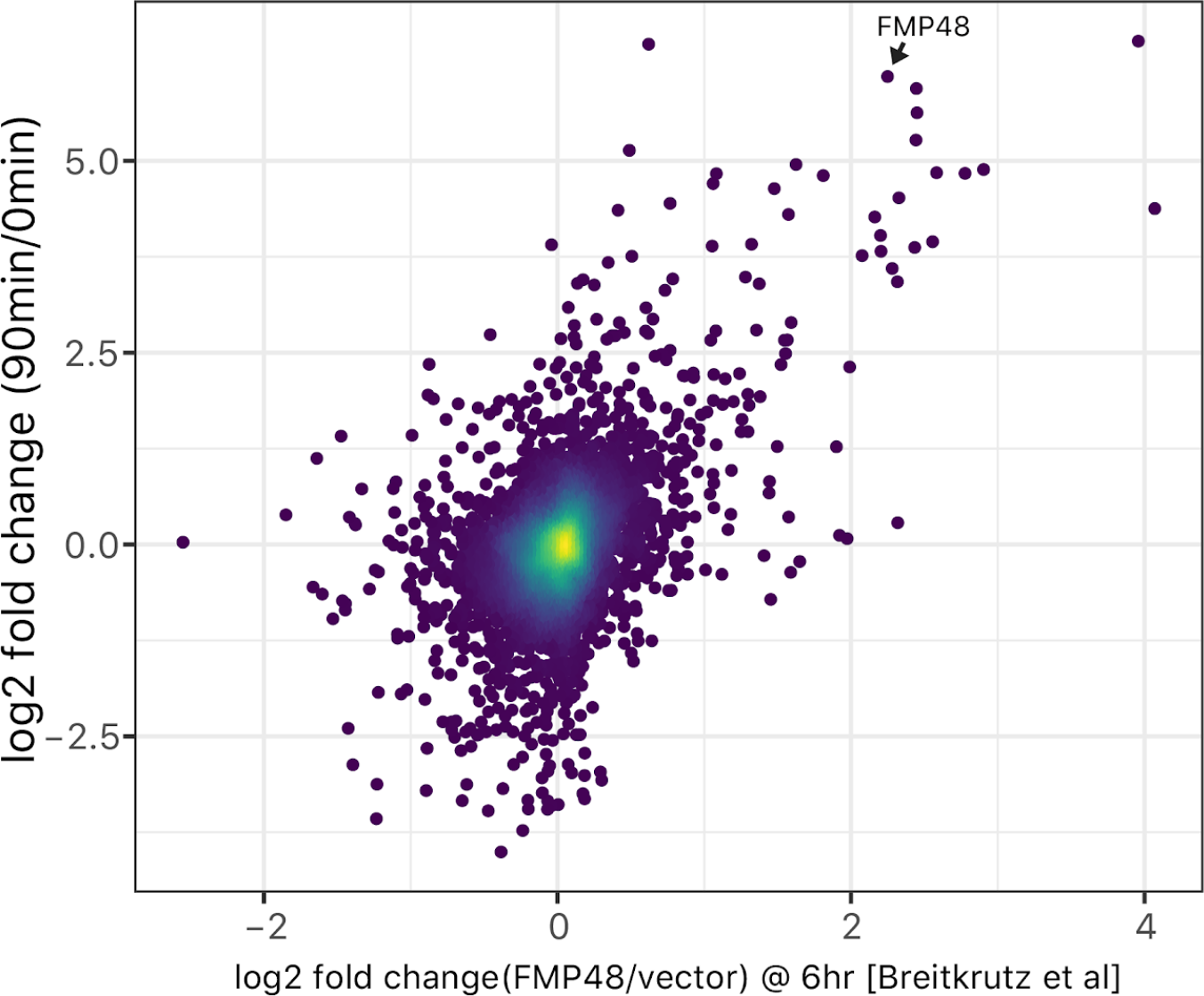
Comparing transcriptome data from two Fmp48 overexpression datasets. Data shown in Figure 5 are plotted along the y-axis. Replicates from Breitkreutz *et al.* (2009) are averaged and plotted along the x-axis. The Pearson correlation is 0.47 and is significant with p-value < 2.2e-16.

## Notes

#### Summary of Updates

minor edits to abstract

http://pin.research.calicolabs.com

## References

1. Ronen M, Botstein D. Transcriptional response of steady-state yeast cultures to transient perturbations in carbon source. Proc Natl Acad Sci U S A. 2006;103: 389–394.

2. Workman CT, Mak HC, McCuine S, Tagne J-B, Agarwal M, Ozier O, et al. A systems approach to mapping DNA damage response pathways. Science. 2006;312: 1054–1059.

3. Simon I, Barnett J, Hannett N, Harbison CT, Rinaldi NJ, Volkert TL, et al. Serial regulation of transcriptional regulators in the yeast cell cycle. Cell. 2001;106: 697–708.

4. Kim H, Shin J, Kim E, Kim H, Hwang S, Shim JE, et al. YeastNet v3: a public database of data-specific and integrated functional gene networks for Saccharomyces cerevisiae. Nucleic Acids Res. 2014;42: D731–6.

5. Harbison CT, Gordon DB, Lee TI, Rinaldi NJ, Macisaac KD, Danford TW, et al. Transcriptional regulatory code of a eukaryotic genome. Nature. 2004;431: 99–104.

6. Lickwar CR, Mueller F, Hanlon SE, McNally JG, Lieb JD. Genome-wide protein-DNA binding dynamics suggest a molecular clutch for transcription factor function. Nature. 2012;484: 251–255.

7. Teytelman L, Thurtle DM, Rine J, van Oudenaarden A. Highly expressed loci are vulnerable to misleading ChIP localization of multiple unrelated proteins. Proc Natl Acad Sci U S A. 2013;110: 18602–18607.

8. Costanzo M, Baryshnikova A, Bellay J, Kim Y, Spear ED, Sevier CS, et al. The genetic landscape of a cell. Science. 2010;327: 425–431.

9. Carlson MRJ, Zhang B, Fang Z, Mischel PS, Horvath S, Nelson SF. Gene connectivity, function, and sequence conservation: predictions from modular yeast co-expression networks. BMC Genomics. 2006;7: 40.

10. Kang Y, Patel N, Shively C, Recio PS, Chen X, Wranik BJ, Kim G, Mitra R, McIsaac RS, Brent MR. Mapping transcription factor networks by comparing TF binding locations to TF perturbation responses. biorxiv. 2019;

11. McIsaac RS, Petti AA, Bussemaker HJ, Botstein D. Perturbation-based analysis and modeling of combinatorial regulation in the yeast sulfur assimilation pathway. Mol Biol Cell. 2012;23: 2993–3007.

12. Greenfield A, Hafemeister C, Bonneau R. Robust data-driven incorporation of prior knowledge into the inference of dynamic regulatory networks. Bioinformatics. 2013;29: 1060–1067.

13. Ronen M, Rosenberg R, Shraiman BI, Alon U. Assigning numbers to the arrows: parameterizing a gene regulation network by using accurate expression kinetics. Proc Natl Acad Sci U S A. 2002;99: 10555–10560.

14. McIsaac RS, Oakes BL, Wang X, Dummit KA, Botstein D, Noyes MB. Synthetic gene expression perturbation systems with rapid, tunable, single-gene specificity in yeast. Nucleic Acids Res. 2013;41: e57.

15. McIsaac RS, Gibney PA, Chandran SS, Benjamin KR, Botstein D. Synthetic biology tools for programming gene expression without nutritional perturbations in Saccharomyces cerevisiae. Nucleic Acids Res. 2014;42: e48.

16. McIsaac RS, Silverman SJ, McClean MN, Gibney PA, Macinskas J, Hickman MJ, et al. Fast-acting and nearly gratuitous induction of gene expression and protein depletion in Saccharomyces cerevisiae. Mol Biol Cell. 2011;22: 4447–4459.

17. Gasch AP, Spellman PT, Kao CM, Carmel-Harel O, Eisen MB, Storz G, et al. Genomic expression programs in the response of yeast cells to environmental changes. Mol Biol Cell. 2000;11: 4241–4257.

18. Teixeira MC, Monteiro PT, Palma M, Costa C, Godinho CP, Pais P, et al. YEASTRACT: an upgraded database for the analysis of transcription regulatory networks in Saccharomyces cerevisiae. Nucleic Acids Res. 2018;46: D348–D353.

19. Chechik G, Oh E, Rando O, Weissman J, Regev A, Koller D. Activity motifs reveal principles of timing in transcriptional control of the yeast metabolic network. Nat Biotechnol. 2008;26: 1251–1259.

20. Song L, Zhang Z, Grasfeder LL, Boyle AP, Giresi PG, Lee B-K, et al. Open chromatin defined by DNaseI and FAIRE identifies regulatory elements that shape cell-type identity. Genome Res. 2011;21: 1757–1767.

21. Arvey A, Agius P, Noble WS, Leslie C. Sequence and chromatin determinants of cell-type–specific transcription factor binding. Genome Res. 2012;22: 1723–1734.

22. Gomes ALC, Wang HH. The Role of Genome Accessibility in Transcription Factor Binding in Bacteria. PLoS Comput Biol. 2016;12: e1004891.

23. Smith SJ, Crowley JH, Parks LW. Transcriptional regulation by ergosterol in the yeast Saccharomyces cerevisiae. Mol Cell Biol. American Society for Microbiology Journals; 1996;16: 5427–5432.

24. Protchenko O, Philpott CC. Regulation of intracellular heme levels by HMX1, a homologue of heme oxygenase, in Saccharomyces cerevisiae. J Biol Chem. 2003;278: 36582–36587.

25. Cherry J. SGD: Saccharomyces Genome Database [Internet]. Nucleic Acids Research. 1998. pp. 73–79. doi:10.1093/nar/26.1.73

26. Zhu C, Byers KJRP, McCord RP, Shi Z, Berger MF, Newburger DE, et al. High-resolution DNA-binding specificity analysis of yeast transcription factors. Genome Res. 2009;19: 556–566.

27. O’Duibhir E, Lijnzaad P, Benschop JJ, Lenstra TL, van Leenen D, Groot Koerkamp MJA, et al. Cell cycle population effects in perturbation studies. Mol Syst Biol. 2014;10: 732.

28. Breitkreutz A, Choi H, Sharom JR, Boucher L, Neduva V, Larsen B, et al. A global protein kinase and phosphatase interaction network in yeast. Science. 2010;328: 1043–1046.

29. Tkach JM, Yimit A, Lee AY, Riffle M, Costanzo M, Jaschob D, et al. Dissecting DNA damage response pathways by analysing protein localization and abundance changes during DNA replication stress. Nat Cell Biol. 2012;14: 966–976.

30. Hackett SR, Zanotelli VRT, Xu W, Goya J, Park JO, Perlman DH, et al. Systems-level analysis of mechanisms regulating yeast metabolic flux. Science. 2016;354. doi:10.1126/science.aaf2786

31. Buchler NE, Gerland U, Hwa T. On schemes of combinatorial transcription logic. Proc Natl Acad Sci U S A. 2003;100: 5136–5141.

32. Chong YT, Koh JLY, Friesen H, Duffy SK, Cox MJ, Moses A, et al. Yeast Proteome Dynamics from Single Cell Imaging and Automated Analysis. Cell. 2015;161: 1413–1424.

33. Bintu L, Buchler NE, Garcia HG, Gerland U, Hwa T, Kondev J, et al. Transcriptional regulation by the numbers: models [Internet]. Current Opinion in Genetics & Development. 2005. pp. 116–124. doi:10.1016/j.gde.2005.02.007

34. Tan M, Luo H, Lee S, Jin F, Yang JS, Montellier E, et al. Identification of 67 histone marks and histone lysine crotonylation as a new type of histone modification. Cell. 2011;146: 1016–1028.

35. Meier L, Van De Geer S, Bühlmann P. The group lasso for logistic regression. J R Stat Soc Series B Stat Methodol. 2008;70: 53–71.

36. Kelley DR, Snoek J, Rinn JL. Basset: learning the regulatory code of the accessible genome with deep convolutional neural networks. Genome Res. 2016;26: 990–999.

37. Klumpp S, Zhang Z, Hwa T. Growth rate-dependent global effects on gene expression in bacteria. Cell. 2009;139: 1366–1375.

