## Supplementary Text for "Time-resolved genome-scale profiling reveals a causal expression network"

### Contents


**1 Growth Conditions** **3**

**2 TF Strain Construction** **3**

**3 RNA extraction, labeling, and hybridization** **3**

**4 Initial Data Processing** **3**

**5 Obtaining the *raw* gene expression dataset** **3**

**6 Obtaining the *cleaned* gene expression dataset** **4**

**7 Obtaining the *noise-value thresholded* gene expression dataset** **4**

**8 Modeling timecourses with sigmoidal or impulse-like dynamics** **5**

**9 Obtaining the *shrunken* gene expression dataset** **6**

**10 Obtaining kinetic parameters for timecourses** **6**

**11 De novo motif discovery** **7**

**12 Modeling Summary** **8**

**13 Dynamical systems modeling** **8**

**14 Linear Regression** **9**

**15 BIC regularization** **10**

**16 Hyperparameters** **11**

**17 Model Validation by Holdout** **11**

**18 Marginal attribution analysis** **12**

**19 Validation Experiments** **14**

### 1 Growth Conditions

For all timecourse experiments, cells were grown under continuous culturing conditions in 500 mL vessels as previously described with minor adjustments [1]. Cultures were aerated with 6 L/min of humidified air at 30°C, maintained at 300 mL, and stirred with a magnetic impeller at 400 RPM. For the majority of experiments, cultures were maintained with minimal medium under phosphate limitation (20 mg/L). Where noted, cultures were maintained under methionine limitation (7.5 mg/L) or under nitrogen limitation (40 mg/L ammonium sulfate). Data for TF experiments performed in methionine limitation were originally published in [2]. Growth rates were maintained from 0.15-0.17 $h^{-1}$. Batch growth in the chemostat vessels was initiated from a 1:60 dilution of a saturated overnight culture prior to turning on the chemostat pumps. Cells were grown to steady state, as determined by culture density, prior to the addition of 1 µM beta-estradiol to the culture and subsequent sampling. Chemostat experiments were performed with either the Infors Sixfors or Multifors systems.

### 2 TF Strain Construction

Parent strains were engineered to constitutively express an artificial transcription factor that is inducible with estradiol. The majority of parent strains used for gene expression analysis contain either GEV or the $Z_3$EV transcription factor. A synthetic promoter fused to the KanMX was PCR amplified and introduced into parent strains via homologous recombination using a standard lithium acetate transformation procedure. Clones containing the synthetic promoter were selected for on rich medium (YPD [1% yeast extract, 2% bacto-peptone, and 2% dextrose] containing G418 (200-300 µg/mL). Primers were designed using custom software in R such that the KanMX-Promoter cassette was introduced directly between a target gene's first methionine residue and its native promoter to prevent the removal of any genomic DNA.

### 3 RNA extraction, labeling, and hybridization

Crude RNA was extracted using a standard acid-phenol procedure. RNA was then purified using either the QIAGEN RNeasy kit or RNAClean Ampure XP beads. 200 ng of cleaned RNA was used as input to generate dye-labeled cRNA using the Agilent Quick-Amp Labeling Kit. Labeled cRNA was cleaned using RNAClean Ampure XP beads. Reference RNA was extracted from DBY12001, a laboratory wild-type strain, grown to steady state in a phosphate-limited chemostat at D = 0.18 $h^{-1}$. Reference and sample RNA were labeled with Cy3-CTP and Cy5-CTP, respectively. Labeled RNA was hybridized to Agilent 8x15k microarrays, which were then washed, scanned, and processed using the Agilent Feature Extraction software with default settings and loess dye bias correction.

### 4 Initial Data Processing

We performed a number of signal processing steps to our microarray datasets as described below. In increasing order of processing, we refer to these as the "raw" dataset, "cleaned" dataset, "noise-model thresholded" dataset, and the "shrunken" dataset.

### 5 Obtaining the *raw* gene expression dataset

The raw measurements for this work consist of one microarray per time point in each timecourse. Each microarray has some number of spots (usually 2) for each gene. All microarrays for the GEV experiments had identical probes, likewise for the ZEV experiments. For each spot, we computed ratio = max(red, C) / max(green, C), where C=2 in arbitrary units. This minimum value application only affects 0.3% of spots. Typical genes have red and green channel measurements above 200. These (red, green, ratio) values for each spot serve as the "raw" data.

### 6 Obtaining the *cleaned* gene expression dataset

For each dataset, we aggregate the data across individual spots. Specifically, for each timecourse, time point, and gene, we aggregated the spot values and measured the minimum value, maximum value, median value, and standard deviation of the values. In the usual case of 2 spots, the median value is equivalent to mean and standard deviation is equivalent to (max-min)/sqrt(2).

At this stage we corrected the most extreme outlier observations. First, for a given sample, we examined the case where, for a given gene, the ratio of the spot values, is larger than four. Since the spots values do not agree, we interpolate the value with the geometric mean across bracketing time points (or neighboring time point in the case of first or last time point). The second class of outliers is where the median ratio moves by a factor of at least four between two time points, and then by a factor of four in the opposite direction for the next time point. In these cases we again replace the central point with the geometric mean of the bracketing points. These corrections apply to less than 0.2% of the data.

In processing gene expression microarrays, crosstalk between red and green channels can occur. When the red channel fluorescence is much larger than the green channel fluorescence, there can be leakage of signal, and the green channel measurement is affected. To identify instances of this occurring, we first computed the green channel ratio relative to the time zero measurement. We then measured the 30% quantile of this value within each timecourse (for time points after t = 0), as well as the 30% quantile of the log-ratio. We then flagged timecourses where the green ratio quantile exceeded a factor of eight, and the log-ratio quantile exceeds a two-fold change. These represent cases where the green channel is increasing a great deal when it should be constant. For these rare cases, we repaired the red to green ratios by duplicating the time zero green channel across the full timecourse. This affects about two dozen TF-gene timecourses (out of more than a million), crucially including GAT3 in the GAT3 induction timecourse.

The GEV and ZEV systems both have characteristic gene expression signatures, including a mild stress response. Previously, it has been shown that Singular Value Decomposition (SVD) is one way to remove such signals [2]. Here, we use a slightly different approach. First, we computed the median time series for each gene in each class of experiments (GEV and ZEV). We then subtracted this median time series, leaving a normalized log-ratio. Because a given gene is not directly or indirectly affected by a transcription factor in most experiments, the median is an accurate reflection of any nuisance time-dependent behavior. We refer to the dataset where outliers are removed and the GEV/ZEV signal is removed as the "cleaned" dataset.

### 7 Obtaining the *noise-value thresholded* gene expression dataset

Biological and technical variability limit the precision of any measurement of gene expression. Under the assumption that any gene is not affected in most experiments, the width of the central quintile (20%) of "cleaned" log-ratio measurements is an accurate estimate of the total noise level for that gene. We assumed log-normal distributions for each gene, mapping central quintile to standard deviations (central quintile = $\pm 0.253\sigma$ for normal). The typical standard deviation (i.e., noise estimate) for cleaned log-ratios (base 2) is approximately 0.1 for genes with green medians above about 200. Genes with lower green medians have higher noise levels, up to about 0.75 (a factor of 1.7).

We then compared this gene-level noise estimate with the typical (median) spot-to-spot error for the genes. We found that the two are mostly uncorrelated, with the spot-to-spot error being significantly smaller. For genes with low green median, the spot-to-spot error accounts for most of the gene error.

Given a noise model, we can now consider which data points represent credible discoveries. We first considered each gene separately, and compare the cleaned log-ratio to the "universal threshold", as defined in [3]. This approach uses the fact that the maximum of N draws from a normal distribution is close to $\sqrt{2 \ln N}\sigma$. If the bulk of the noise is Gaussian, we can control the probability of at least one false positive. For the 1,500 non-zero time points for a given gene, this corresponds to $3.8\sigma$. We flagged all measurements that exceeded this threshold in either direction (a total of 3.3% of the time points). The first level of thresholding is to set all timecourses with no points passing threshold to exactly zero.

Noise levels can also vary between microarrays. Thus, we zeroed out any timecourse that might have signal according to the gene-only noise model, and then computed the microarray-level noise from the central quintile of the genes that were not filtered out. We then constructed a simple metric to describe undesirable timecourses. Timecourses with a single significant detection (not at the final time point), or timecourses with non-consecutive significant detections, are not likely to be true positives. We then chose a weighting factor to minimize the fraction of detected timecourses that are of these types [noise model: $\sigma^2_{total} = (5/6)\sigma^2_{gene} + (1/6)\sigma^2_{microarray}$]. This removed obvious microarray-level artifacts while preserving the signal in timecourses where a large fraction of the genome is affected.

Collectively, the full cleaned dataset is 1,693 microarrays in 217 timecourses, each with 6,175 genes. Of the 1.34 million time series, 118,134 (8.8%) have at least one point that passes the full noise model. 60,968 (4.5%) have multiple points that pass the noise model. Of the time series with only single detection, about half occur in the final time point, suggesting that these are genuine late changes. Conversely, 9,329 time series possessed gaps between detections suggesting that their changes may be non-biological. These two filters (removing non-final singletons and non-consecutive changes) left 79,709 timecourses (5.9%), which were used to train the whole-cell dynamical systems model.

### 8 Modeling timecourses with sigmoidal or impulse-like dynamics

Next, we wanted to identify timecourses that most likely contain smooth, biologically-feasible dynamics. In order to capture dynamics without overly restricting the types of dynamics that might exist, each timecourse with observation-level signal was fit with the phenomonological impulse model of Chechik and Koller [4]. This model expresses timecourses as the sum of two sigmoidal changes each which have a characteristic amplitude and time constant. This impulse model has been sufficient to capture complex pattern in expression and metabolomic data. timecourses were considered well fit by an impulse model if over 80% of expression variation could be accounted for through the impulse fit (i.e., $\sum(\hat{f}_{ij} - f_{ij})^2 / \sum f_{ij}^2 < 0.2$).

This filter removed highly inconsistent responses, but in some cases dynamics were clearly driven by an outlier yet an impulse model still fit the data well. To remove these cases, two heuristics were applied to separately remove outliers driven by the time zero measurement, and those occurring at an intermediate time point. In the first few minutes of an induction experiment, few strong transcriptional changes exist besides the induced transcription factor. In practice if the time zero point was noisy, measurements that are normalized together with respect to this value will be systematically higher or lower than expected, resulting in sharp early change in expression. Such changes were removed by filtering timecourses where the largest absolute change in expression was between the first and second timepoint and in which the impulse model explained less than 98% of variation is explained by the impulse model (the latter filter recovered early, strong changes such as the primary induction event). The second case of pathological timecourses were cases where a single outlier measurement (beyond what could be detected by our Gaussian noise model) was not appropriately reflected in either the preceding $f_{t-1}$, nor the succeeding $f_{t+1}$ time point despite fine temporal samples. Large changes followed by a return to baseline were distinguished from continuous timecourse responses by removing timecourses with a serial sum of squares to the total sum of squares ratio of less than 1.25 (i.e., $\frac{\sum_t^{T-1}(f_{t+1}-f_t)^2}{\sum_t (f_t)^2} < 1.25$).

Starting with the 118,134 timecourses that had at least one point that passed the full noise model, we filtered - based on the timecourse-level patterns - 18,098 timecourses from further consideration, leaving 100,036 timecourses which contain clear impulse-like or sigmoidal dynamics.

### 9 Obtaining the *shrunken* gene expression dataset

We believe that each of the previously mentioned 100,036 timecourses contains timecourse-level signal, but not necessarily at every time point. Indeed changes in the first 5 minutes are very rare (aside from the induced transcription factor), while near the end of the experiment, the vast majority of these timecourses will be different than their pre-induction expression level. To avoid the case of inappropriately interpreting early, often very weak expression variation as signal, we want to shrink signals to zero to the extent that they are consistent with the estimated noise-level of the gene $\hat{\sigma}_{ij}^2$. To determine whether each observation in a timecourse is consistent with the noise model, we perform a Wald test for every observation ($W_{ijt} = f_{ijt}/\hat{\sigma}_{ijt}$). As expected, Wald p-values at early timepoints are more uniformly distributed while later timepoints skew very strongly towards small p-values. This difference in the fraction of null hypotheses at a given time point ($\pi_0$) can be modeled as a monotonically decreasing function of time $\pi_0(t)$ using the functional false discovery rate [5, 6]. In order to make use of this approach for shrinkage estimation, we consider that each observation can be thought of as a mixture of two states, a null state where $\hat{f}_{ijt} = 0$ and an alternative state reflecting our experimental measurement: $\hat{f}_{ijt} = f_{ijt}$, weighted based on the relative support for an observation belonging to each state ($w_{ijt}$):

$$f_{ijt}^{\text{shrunken}} = w_{ijt} \times 0 + (1 - w_{ijt}) f_{ijt}^{\text{cleaned}}$$

$w_{ijt}$ can be estimated across all the observation of a single timepoint (using the above estimated $\pi^0(t)$) using the local false discovery rate (LFDR) [6]. The LFDR is an estimate of the FDR of a single observation. Because $\pi_0$ is smaller for later timepoints, observations later in timecourses tend to be more similar to their measurements, while early timepoints are shrunk more aggressively towards zero. This is the "shrunken" dataset.

### 10 Obtaining kinetic parameters for timecourses

To compare the behavior of timecourses it is useful to be able to summarize both the timing and strength of transcriptional changes. To capture such simple dynamics, a sigmoidal model may often be sufficient:

$$y(t) = v_{\text{inter}} \frac{1}{(1 + \exp(-\text{rate} * (\text{time} - t_{\text{rise}})))}$$

However, in some cases the impulse model utilized above to define feasible timecourse-level signal may be necessary to capture complex dynamics:

$$y(t) = \frac{(v_{\text{final}} + (v_{\text{inter}} - v_{\text{final}}))}{1 + \exp(-\text{rate} * (\text{time} - t_{\text{rise}}))} * \frac{1}{1 + \exp(\text{rate} * (\text{time} - t_{\text{fall}}))}$$

In order to make use of kinetic information, we would like to determine when a sigmoidal versus an impulse model is supported and when and how strongly activations/inhibitions occur.

While in the previous section we utilize the impulse model to define whether a timecourse can be reasonably described by an impulse relationship, here we are interested in generating interpretable parameters from such fits. Since sigmoidal or impulse models can fit the data in peculiar ways that are not biologically feasible (e.g., negative rate coefficients, impulses which fall before they rise, very strong late activations which are similarly captured with weaker activations), often with a similar fit to a reasonable parameterization, some constraints on parameters, through the use of priors, can greatly improve interpretability. To adapt this problem from non-linear least squares problem to one where we could apply prior constraints, sigmoid/impulse model parameters were estimated as maximum posterior (MAP) estimates ($\arg\max_{\Omega} \Pr(\Omega|\mathbf{y})$) over 100 initializations.

Gaussian likelihood defined the departures between observed fold changes and the non-linear prediction of an impulse or sigmoid model, while the parameters ($\Omega$) of these predictions ($\hat{y}_i(t_i, \Omega)$) were constrained with Gaussian and Gamma priors:

$$
\begin{aligned}
\Pr(\Omega|\mathbf{y}) &\propto \Pr(\mathbf{y}|\Omega)\cdot\Pr(\Omega)\\
\Pr(\mathbf{y}|\Omega) &= \prod_{i=1}^{I}\mathcal{N}\left(\hat{y}_i(t_i,\Omega);\mu=y_i,\sigma=\hat{\sigma}_{ijt}\right)\\
\Pr(\Omega^{\text{sigmoid}}) &= \mathcal{N}(v_{\text{inter}};0,1)\cdot\Gamma(\beta;2,0.25)\cdot\Gamma(t_{\text{rise}};2,25) \quad \text{sigmoid}\\
\Pr(\Omega^{\text{impulse}}) &= \mathcal{N}(v_{\text{inter}};0,1)\cdot\Gamma(\beta;2,0.25)\cdot\Gamma(t_{\text{rise}};2,25)\cdot\\
&\quad \mathcal{N}(v_{\text{final}};0,1)\cdot\Gamma(t_{\text{fall}}-t_{\text{rise}};2,25) \quad \text{impulse}
\end{aligned}
$$

Each timecourse was fit with both a sigmoidal and an impulse model and we sought to determine which model best fit each timecourse. Since the sigmoidal model is a simpler, nested version of the impulse model (with $t_{\text{fall}}=\infty$), the likelihood ratio test was used to determine whether the sigmoid was significantly improved by the two extra parameters of the impulse model:

$$\log\Pr(\mathbf{y}|\Omega^{\text{impulse}})-\log\Pr(\mathbf{y}|\Omega^{\text{sigmoid}})\sim\chi^2_2$$

For $1,785$ timecourses, the impulse model fit significantly better than the sigmoid at Benjamini-Hochberg-based FDR of 0.001. These timecourses were said to have impulse dynamics while the remaining $98,251$ timecourses exhibited sigmoidal dynamics.

### 11 De novo motif discovery

To construct data-driven binding motifs for each induced transcription factor and suggest other transcription factors that may be operating in each induction experiment, we sought to identify cis-regulatory motifs which are enriched in the promoters of regulated genes and, where possible, attribute a known regulator to those motifs. To identify enriched motifs, we utilized the regular-expression-based software, DREME [7], to identify short (8-mers or shorter) motifs that are enriched in a set of primary sequences relative to a set of control sequences. The sequences we focused on for all motif analyses were the promoters of yeast genes (defined as 500 basepairs upstream of each gene; downloaded from Ensembl on 2018-01-03). To identify regulatory motifs, for each experiment, sequences in the promoters of regulated genes (sigmoid or impulse responses) were compared to non-differentially expressed control sequences. Similarly, motifs associated with impulse-like behavior were identified by identifying motifs enriched in the promoters of genes with impulse kinetics, using the promoters of genes with sigmoidal kinetics as control sequences. To attribute identities to each motifs, probability weight matrices (PWMs) based on binding data were downloaded from Yeastract [8] and then matched to each DREME motif using TOMTOM [9].

To investigate whether genes containing an enriched motif exhibit stereotypical kinetics, we investigated whether variation in regulatory kinetics across genes could be predicted based on promoter composition. To carry-out such comparisons, the promoters of regulated genes were matched to each identified motif based on enrichment of high-scoring PWM k-mers (Pr(sequence | PWM)) in the promoters of regulated compared to control promoters (using the same primary vs. control comparisons as for DREME). Briefly, this was done by ordering sequences in descending order of PWM score match and then using a rolling mean estimate of k-mer frequency to estimate the PWM score cutoff where enrichment in the primary sequences decreases to a heuristic 1.5-fold cutoff. Using this approach, each gene responding in a given experiment was summarized based on how many times each motif was detected and what its strongest PWM match was. To determine whether any motif was associated with variation in a kinetic property ($v_{\text{inter}}, t_{\text{rise}}$, rate for regulated genes, with $v_{\text{final}}$ and $t_{\text{fall}}$ added for impulse genes), each kinetic coefficient was regressed on a motif-by-motif basis using ordinary least squares (OLS) on three summaries of each motif's presence. These three predictors were: the top enriched PWM match for the gene, a binary variable indicating whether one or more motifs were present and counts of how many PWM matches were found. OLS t-statistic p-values were separately FDR controlled for each type motif summary ("best match", "motif present", "# of matches") and model type (regulation or impulse) [6].

### 12 Modeling Summary

We pursued a linear regression approach to modeling the dynamical system model of transcription. We constructed an estimator of the time derivative of the gene expression. We treat this as the dependent variable. We then fit a linear model to extract the coefficients of the dynamical system. This works because the time derivatives of the gene expression levels are modeled as linear functions of the gene expression levels, possibly with quadratic terms as well. We note that this does not actually correspond to a full solution of the dynamical system, but merely requiring point-wise consistency with the dynamical system description. Selection of regularization levels with cross validation yielded a model for the transcriptional effects of gene expression levels. This model was interrogated to identify which regulators were most important for predicting observed expression changes in each timecourse.

Ten predicted regulators which lacked an induction experiment were selected for experimental validation. A separate induction timecourse was generated for each predicted regulator, and three regulators had strong downstream transcriptional responses which were also highly enriched for the effects predicted by the model.

### 13 Dynamical systems modeling

We constructed a linear model based on a simple dynamical system model of genome-wide expression evolution. The time rate of change of a gene is modeled as being affected by the expression levels of any of the genes in the genome, possibly linearly, or proportionally to the product of their expression levels. We can transform this equation into the same units as the cleaned data, with values at time zero divided out. We convert this into a regression problem by constructing a derivative estimator from the time series data. We then treat the left-hand side of the equation (i.e., the time derivative) as the dependent variable, modeled by the right-hand side (i.e., the linear and quadratic terms) as the independent variables.

$$\frac{d}{dt}z_i = \sum_j [A_{ij}z_j + B_{ij}z_iz_j] + D_i.$$

This system of equations presumes the natural units and reaction rates for gene expression. We only have the microarray measurements. We absorb the relation between transcript abundance and measured photons into the definitions of a new set of variables and a new set of coefficients and fit the normalized system. Our final variable $y$ is the normalized ratio, y(t)=(r(t)/g(t))/(r(0)/g(0)), and y(t=0)=1 by definition.

$$\frac{d}{dt}y_i = \sum_j [\alpha_{ij}y_j + \beta_{ij}y_iy_j] + \delta_i.$$

The $\alpha$ matrix describes the linear effect of one gene's expression on another, while the $\beta$ matrix describes the same effect when it is also proportional to the target genes expression level. The $\delta$ vector represents a background level of transcription.

When an experiment begins, the yeast is approximately in a steady state. The population average gene expression should therefore be time independent at that point. We can subtract out this behavior explicitly, which would highlight any genes that violate this assumption. This $\sigma$ vector explicitly describes the deviation from steady state at the start of the experiment.

$$\begin{aligned} \sigma_i &= \sum_j [\alpha_{ij} + \beta_{ij}] + \delta_i, \\ \frac{d}{dt}y_i &= \sum_j [\alpha_{ij}(y_j - 1) + \beta_{ij}(y_iy_j - 1)] + \sigma_i. \end{aligned}$$

Linear modeling of the gene expression levels can yield a negative numbers of transcripts which are clearly unphysical. If we instead model the log-ratios, this can never happen. Dividing each side by the time-dependent gene expression level $y_i(t)$ yields a new equation with the log derivative as the left-hand side. This equation is exactly equivalent, but notice that the source $\alpha$, $\beta$ terms and intercept $\sigma$ terms now have the transcript ratio in the denominator.

$$\frac{d}{dt}\ln(y_i) = \sum_j \left[\alpha_{ij}\frac{y_j - 1}{y_i} + \beta_{ij}\frac{y_i y_j - 1}{y_i}\right] + \sigma_i \frac{1}{y_i}.$$

Enforcing the initial steady state means $\sigma_i = 0$, reducing to Eq. 1 in the main text.

We can invert the problem by simply integrating. Now, the left-hand side is the gene expression level, with the time zero subtracted. The right-hand side is the time integral over earlier time points. Thus, the expression level of a gene can be modeled by earlier-time averages of the expression levels of other genes.

$$\ln(y_i(t)) = \int_0^t dt' \left\{ \sum_j \left[\alpha_{ij}\frac{y_j(t') - 1}{y_i(t')} + \beta_{ij}\frac{y_i(t')y_j(t') - 1}{y_i(t')}\right] + \sigma_i \frac{1}{y_i(t')} \right\}.$$

### 14 Linear Regression

Translating the dynamical system described above into the language of linear regression is straightforward. In its simplest form, the dynamics of a single gene in a single intervention experiment is represented by a design matrix with rows corresponding to the (usually 8) time points and columns corresponding to the 6175 genes being measured. We first construct an estimator for the time derivative of the gene in question. This will reduce the effective number of independent microarrays by one per timecourse. N points in a timecourse will provide N-1 derivative estimators. We choose to take the average of the first order forward and backward differences as our estimator, but note that this is not always the symmetric difference since the time samples are not uniform. Furthermore we assume that the backward difference at time zero is zero, and that the forward derivative at the end of the timecourse is also zero. However, we note that the N derivative estimates for the N time points are not linearly independent with these assumptions. Equivalently, if we consider a differencing operator that acts on a timecourse, applying it to a vector of 1s yields all zeros. Thus, the differencing operator is not invertible.

Leaving out the quadratic and intercept terms for clarity, the "derivative" and "integral" models can be written with the following equations, respectively:

$$\sum_Q D^{PQ}\ln(y_i^Q) \sim \sum_j \alpha_{ij}\frac{y_j^P - 1}{y_i^P},$$

$$\ln(y_i^P) \sim \sum_{j,Q} \alpha_{ij} D^{-1,PQ}\frac{y_j^Q - 1}{y_i^Q},$$

where the $P, Q$ indicices now refer to time.

The integral version models the data at a time point as a particular sum over earlier time points. In general, time sampling is not uniform. The majority of timecourses sampled at t = 0, 5, 10, 15, 20, 30, 45, 90 minutes. In this case, the difference and integral operators are, respectively:

$$D = \frac{1}{10 \text{ min}} \begin{pmatrix} -1 & 1 & 0 & 0 & 0 & 0 & 0 & 0 \\ -1 & 0 & 1 & 0 & 0 & 0 & 0 & 0 \\ 0 & -1 & 0 & 1 & 0 & 0 & 0 & 0 \\ 0 & 0 & -1 & 0 & 1 & 0 & 0 & 0 \\ 0 & 0 & 0 & -1 & 1/2 & 1/2 & 0 & 0 \\ 0 & 0 & 0 & 0 & -1/2 & 1/6 & 1/3 & 0 \\ 0 & 0 & 0 & 0 & 0 & -1/3 & 2/9 & 1/9 \\ 0 & 0 & 0 & 0 & 0 & 0 & -1/9 & 1/9 \end{pmatrix}$$

$$D^{-1} = 10 \text{ min} \begin{pmatrix} 0 & 0 & 0 & 0 & 0 & 0 & 0 & 0 \\ 1 & 0 & 0 & 0 & 0 & 0 & 0 & 0 \\ 0 & 1 & 0 & 0 & 0 & 0 & 0 & 0 \\ 1 & 0 & 1 & 0 & 0 & 0 & 0 & 0 \\ 0 & 1 & 0 & 1 & 0 & 0 & 0 & 0 \\ 2 & -1 & 2 & -1 & 2 & 0 & 0 & 0 \\ -1 & 2 & -1 & 2 & -1 & 3 & 0 & 0 \\ 8 & -7 & 8 & -7 & 8 & -6 & 9 & 0 \end{pmatrix}$$

The full regression dataset for the dynamics of the gene in question is constructed by row-wise stacking of the 200+ design matrices corresponding to the intervention experiments. We then repeat this for each of the 6175 genes.

### 15 BIC regularization

The previously explained regression formulation of the dynamic model was fit using lasso regression, and regularization paths were fit using glmnet [10].

Normally the regularization parameter of lasso regression is fit by straightforward cross-validation. Naively, we would use cross validation to select a $\lambda$ for the dynamic model of each gene's dependence on other genes separately. However, there are a number of practical problems that make this inappropriate on this data.

First, although there are around 1600 rows in the design matrix that underlies the regression, these represent only 200 timecourses of 8 time points each. The point being that the interdependency of the data limits the kinds of cross-validation that are meaningful (e.g. ones in which whole time series are left out) and substantially reduces the effective sample size. In practice, separate cross validation for each gene's model is unacceptably unstable.

To explain our alternative approach, we make use of the theory, based on Stein's unbiased risk estimation approach, that the number of nonzero coefficients in lasso regression at a particular choice of $\lambda$ is an unbiased estimate of the degrees of freedom of the model [11].

Accordingly, unbiased estimates of the BIC and AIC criteria can be formulated as $-2\log(L)+c\log(n)\widehat{df}$ where $L$ is the likelihood of the data, $\widehat{df}$ is the number of nonzero coefficients in the model, and $c$ is 1 or $2/\log(n)$, respectively. Note that the dependence of L, $\hat{y}$, $\widehat{df}$, etc., on $\lambda$ is suppressed in the notation, for brevity. Let sse $= ||y - \hat{y}||^2$. Then, for a homoscedastic Gaussian model, the deviance, D, i.e. -2 times the log-likelihood, follows:

$$D = \frac{\text{sse}}{\sigma^2} + n\log(\sigma^2)$$

where sse stands for the sum-squared errors of the model. Since D is minimized at $\hat{\sigma}^2 = \frac{\text{sse}}{n}$, we obtain:

$$\hat{D} = n\left(1 + \log\left(\frac{\text{sse}}{n}\right)\right)$$

This results in the following penalized criterion:

$$\text{criterion}_{v1} = n\left(1 + \log\left(\frac{\text{sse}}{n}\right)\right) + c\log(n)\widehat{df}$$

Rather than apply criterion$_{v1}$ naively, we first address these concerns:

1. To be resilient to the small sample variability of the sse statistic (especially at small values)
2. To compensate for the fact that the homoscedastic noise model does not hold (most importantly because of time-dependence)
3. It is impractical to find a separate regularization parameter for each gene model separately by usual means, because the selections fluctuate too much.

To address the the first point, we can take advantage of the gene expression noise modeling, which gave us estimates of a noise level for each gene, $\tau_g$. In relative units we can write $s^2 = (\text{sse}/n)/\tau_g^2$. Specifically, it is implausible for the predictive model to be able to explain the data substantially better than the $\tau_g$ ($s^2 < 1$) – these are likely to be due to chance.

We can rewrite $\hat{D}$ in terms of $s^2$ as: $\hat{D} = n(1 + \log(\text{sse}/n)) = n(1 + \log(s^2 * \tau_g^2)) = n\log(s^2) + n + 2n\log(\tau_g)$.

The second term is constant with respect to $\lambda$ and may be omitted for model selection purposes. For the $n\log(s^2)$ term, however, by the above argument, this is a suitable functional form when $s^2$ is large, but not when $s^2$ is small. Various arguments can be made, but the form we finalized on was to replace $\log(s^2)$ with $\text{arcsinh}(s^2 - 1)$. This choice preserves the log-like behavior for large $s^2$, and the local linearity near 1, while giving a small maximum reward for the (probably chance) event that $s^2 < 1$. In summary the criterion we use for model selection is:

$$\text{criterion}_{v2} = n\ \text{arcsinh}(s^2 - 1) + c\log(n)\widehat{df}$$

Finally, we addressed points 2 and 3 by letting $c$ be a tunable hyper-parameter whose value is shared across the dynamic expression models of all genes, and whose value will be chosen by a global cross-validation criterion.

### 16 Hyperparameters

For clarity, we enumerate the model choices (hyperparameters) that were evaluated. We started from "cleaned" dataset as described above.

- Data Preprocessing: Thresholding/Filtering/Gene+Chip noise model/LFDR
- Model formulation: Integral versus derivative
- Log vs. Linear: Dependent variable is the log derivative?
- Intercept: Allow the time derivative to be non-zero at t=0?
- Standardize: Covariates are scaled to unit normal before regression?
- Quadratic: Include quadratic independent variables?
- Regularization: Magnitude of the BIC adjustment

Data preprocessing is a discrete choice among thresholding levels. The next five are binary choices. The regularization level is, in principle, chosen from a continuum. In practice, we make a discrete selection among values separated by factors of 2, the factor 1.0 indicating the naive BIC calculation.

### 17 Model Validation by Holdout

Ideally, we would like to do a leave-one-out holdout analysis on the space of hyperparameters, at the transcription factor level. Since there are 200+ distinct transcription factor experiments (a few with replicates), and 6000+ genes, this involves computing 1.2M regression paths per choice of discrete hyperparameters. However, this is very computationally expensive. Therefore, we identified several transcription factors that remain active in the models out-of-sample. These are *CIN5, DAL80, FKH1, GAL4, GRX4, HAC1, HMS1, LEO1, MSN4, RDR1*, and

*UGA3*. For this reduced set we exhaustively searched 128 choices of discrete hyperparameters. There are four binary choices (choosing not to standardize variables) and eight thresholding levels: none, LFDR, (zero, hard, soft thresholding) x (do continuity filtering). Including models with no holdouts, this corresponds to 12x128x6175 = 9.5M regularization paths computed. From these, we selected 34 candidates for full holdout runs (including checking a few where we do standardize variables). Over all hyperparameter and holdout selections, over 8000 regularization paths were generated for each transcript, over 50M in total.

To score a fully specified model (i.e. a complete specification of model hyperparameters) we used the method described in Algorithm 1. In order to score a holdout experiment, the model coefficients were interpreted as predictions about which genes would move up by 2-fold or down by 2-fold.

In ordinary cross validation, one tries to minimize out-of-sample squared-error. Instead, we choose to maximize out-of-sample coefficient correctness (i.e., the coefficients' ability to make out-of-sample predictions about observed 2-fold changes). The models that scored best (on average) making out-of-sample predictions, including consideration of the sign (induction vs. repression), were called "best".

According to these criteria, the best performing model was:

- Data Preprocessing: zero thresholding, with continuity filter
- Derivative: dependent variable is the time derivative estimate
- Log: dependent variable is the log derivative
- No Intercept: prediction is forced to zero at time zero
- No Standardize: covariates are not whitened before regression
- Quadratic: quadratic terms are included in the design
- Regularization: BIC adjustment = 0.5

With these hyperparameters, we construct a final model with no experiments held out. The model takes the form of a number of coefficients stating that gene A induces or suppresses gene B.

### 18 Marginal attribution analysis

The whole-cell regression model readily provides two important summaries of regulation. First, the estimated coefficients of the regression model ($\beta$) capture regulatory potential ($\beta = \partial y/\partial x$). Second, the regression model combines regulatory potential with variation in regulators through observation-level fitted values ($X\beta$) to summarize how regulation unfolded within an experiment. To interpret regulation, we want to be able to identify the major variable regulators underlying regulatory phenomena of interest such as the impulse-like dynamics of Aft1 or variable timing of expression induction or repression. Since realized regulation is a property of an experiment, the fitted model ($X\beta$) informs whether the model collectively predicts regulatory expression changes of interest and marginal interpretation of components of this model ($x^T\beta$) can be used to attribute regulation to specific regulators.

To attribute regulation, we first identify instances of regulation that are reasonably predicted by the whole-cell model. Each instance of realized regulation is a change in expression occurring over a period of time. These transitions in the data can be readily understood within the framework of the previously discussed parametric models since these models indicate which regulatory phenomena to track and the saturation of the sigmoidal curves implies the period of time over which regulation is unfolding:

$$t\{\text{sat} = x\} = t_{\text{coef}} + \log(x/(1-x))/\beta$$

Here, $t\{\text{sat} = x\}$ is the time at which the sigmoid is $x$ saturated (i.e., 90% of the response having occurred equates to $x = 0.9$). $t_{\text{coef}}$ is the half-max time coefficient of the transition ($t_{\text{rise}}$ or $t_{\text{fall}}$ for a given phenomena). With this convention, the end-points of each rise and fall phenomena, $t_{\text{start}}, t_{\text{end}}$, were defined as the time which the rise or fall sigmoid was 5% to 95% saturated.

**Algorithm 1** Hyperparameter search for dynamical systems modeling.

**Require:** raw data $\mathbf{D}_{\text{RAW}}$ $\triangleright$ (experiment, time) X gene
**Require:** gene set $G = \{g_i\}$ $\triangleright$ 6175 genes
**Require:** experiment set $H = \{h_j\}$ $\triangleright$ 200+ genes with direct experiments
**Ensure:** $H \subset G$
**Require:** hyperparameter sets $\Theta = \{\theta_k\}$ $\triangleright$ each $\theta_k$ is a collection of values
**Require:** BIC correction factors $\vec{B}$

**for** $\theta_k \in \Theta$ **do**
$\quad \mathbf{D} \leftarrow \textsc{PreprocessData}(\mathbf{D}_{\text{RAW}}, \theta_k)$
$\quad$ **for** $h_j \in H$ **do**
$\quad\quad \mathbf{D}_H \leftarrow \textsc{HoldOutExperiment}(\mathbf{D}, h_j)$
$\quad\quad \{\vec{\beta}_i(\vec{B})\} \leftarrow \textsc{FitModel}(\mathbf{D}_H, \theta_k, \vec{B})$
$\quad\quad \mathbf{D}_K \leftarrow \textsc{KeepOnlyExperiment}(\mathbf{D}, h_j)$
$\quad\quad S_j(\vec{B}) \leftarrow \textsc{ScoreModelCoefficients}(\{\vec{\beta}_i(\vec{B})\}, \mathbf{D}_K)$
$\quad$ **end for**
$\quad \Omega(\theta_k, \vec{B}) \leftarrow \textsc{CollateHoldoutScores}(\{S_j(\vec{B})\})$
**end for**
$\theta, B \leftarrow \text{argmax } \Omega(\Theta, \vec{B})$
$\mathbf{D} \leftarrow \textsc{PreprocessData}(\mathbf{D}_{\text{RAW}}, \theta)$
$\{\vec{\beta}_i^{\text{FINAL}}\} \leftarrow \textsc{FitModel}(\mathbf{D}, \theta, B)$

**procedure** $\textsc{FitModel}(\mathbf{D}_{\text{in}}, \theta_k, \vec{B})$
$\quad$ **for** $g_i \in G$ **do**
$\quad\quad$ **if** $g_i \in H$ **then**
$\quad\quad\quad \mathbf{D} \leftarrow \textsc{HoldOutExperiment}(\mathbf{D}_{\text{in}}, g_i)$ $\triangleright$ $g_i$ experiment held out
$\quad\quad$ **else**
$\quad\quad\quad \mathbf{D} \leftarrow \mathbf{D}_{\text{in}}$
$\quad\quad$ **end if**
$\quad\quad \mathbf{X} \leftarrow \textsc{ConstructDesignMatrix}(\mathbf{D}, g_i, \theta_k)$
$\quad\quad \vec{y} \leftarrow \textsc{ConstructDependentVariable}(\mathbf{D}, g_i, \theta_k)$
$\quad\quad \vec{\beta}_i(\vec{\lambda}) \leftarrow \textsc{FitLASSO}(\mathbf{X}, \vec{y}, \theta_k)$ $\triangleright$ model is $\vec{y} \sim \mathbf{X}\vec{\beta}$
$\quad\quad \vec{\beta}_i(\vec{B}) \leftarrow \text{BIC\_}\textsc{CorrectionFactor}(\mathbf{X}, \vec{y}, \vec{\beta}_i(\vec{\lambda}))$ $\triangleright$ $\vec{\lambda} \rightarrow$ BIC
$\quad$ **end for**
$\quad$ **return** $\{\vec{\beta}_i(\vec{B})\}$
**end procedure**

To determine when the whole-cell model accounts for an appreciable fraction of observed regulatory changes, model-predicted fold-changes over each regulatory interval $f^{\text{model}} = \log 2(y[t_{\text{end}}]) - \log 2(y[t_{\text{start}}])$ were compared with the observed change in that gene's expression over the regulatory interval $f^{\text{observed}} = \log 2(x[t_{\text{end}}]) - \log 2(x[t_{\text{start}}])$. Since $t_{\text{start}}, t_{\text{end}}$ will generally occur between time-points linear interpolation of both $\log 2(y)$ and $\log 2(x)$ using the two closest timepoint was used to infer these intermediate expression states. For $47,802$ responses (rises or falls), the model had some predictive value based on the following cutoff:

$$\min([f^{\text{model}}, f^{\text{observed}}])/\max(f^{\text{model}}, f^{\text{observed}}) > 0.2$$

Marginal attribution analysis was used to dissect total model fits into the marginal contributions of each regulator. Here the marginal attribution of each regulator to the a response was defined as:

$$\psi_{ijkz} = |f^{\text{model}}_{ijkz}| / \sum |f^{\text{model}}_{ikz}|$$

Here, $\psi_{ijkz}$ is the model's predicted proportional control of a regulator $j$ to a gene $i$ in experiment $k$ for the $zth$ response (rise or fall) occurring in that timecourse. $\psi$'s (filtered to $\psi > 0.2$) were used to interpret the major regulator(s) contributing to observed responses in individual experiments and in the meta-graph of cross-experiment regulation.

### 19 Validation Experiments

Based on our modeling results, ten predicted latent regulators were chosen for experimental validation. Selected regulators fell into three classes of predictions:

- Regulators which were predicted to drive the strong impulse behavior of Pho4 induction experiment: Pho5, Pho11, Phm6
- Regulators predicted to affect many targets spanning all experiments: **Hmx1**, Arn2, Anb1, YGL117W
- Regulators predicted as hubs, operating in many experiments: **Stp4**, **Fmp48**, YGR066C

Each predicted regulator was separately induced and genome-wide expression was tracked at eight time points using the same experimental and computational methodology used to construct the "cleaned" dataset. Predicted regulators whose effects were primarily restricted to genes involved in the non-specific induction stress response (Phm6) were removed from consideration since their effects are either due to or confounded with the stress response. Predicted regulators were also removed from consideration if the induced gene increased by less than four-fold during the experiment (Pho11). For the remaining eight validation experiments, a regulator's predicted targets were compared to experimental changes (absolute fold-change $> 0.5$ at any time point) by constructing a contingency table and applying a $\chi^2$-test to evaluate the independence of the marginal effects.

### References


[1] Saldanha AJ, Brauer MJ, Botstein D. *Nutritional Homeostasis in Batch and Steady-State Culture of Yeast.* Mol Biol Cell. 2004 Sep;15(9)4089-103.

[2] McIsaac RS, Petti AA, Bussemaker HJ, Botstein D. *Perturbation-based analysis and modeling of combinatorial regulation in the yeast sulfur assimilation pathway.* Mol Biol Cell. 2012 Aug;23(15):2993-3007.

[3] Donoho DL, Johnstone IM, Kerkyacharian G, Picard D. *Wavelet shrinkage: asymptopia?* Journal of the Royal Statistical Society Series B. 1995; 57(2):301-369.

[4] Chechik G, Koller D. *Timing of gene expression responses to environmental changes.* J Comput Biol. 2009 Feb;16(2):279-90.

[5] Chen X, Robinson DG, Storey JD *The Functional False Discovery Rate with Applications to Genomics.* bioRxiv 241133; doi: https://doi.org/10.1101/241133.

[6] Storey JD, Tibshirani R. *Statistical significance for genomewide studies.* PNAS. 2003. 100(16):9440-5.

[7] Blainey TL. *DREME: Motif discovery in transcription factor ChIP-seq data.* Bioinformatics. 2011. 27(12):1653-1659.

[8] Teixeira MC, Monteiro PT, Palma M, Costa C, Godinho CP, Pais P, Cavalheiro M, Antunes M, Lemos A, Pedreira T, Sa-Correia I. *YEASTRACT, an upgraded database for the analysis of transcription regulatory networks in Saccharomyces cerevisiae* Nucleic Acids Research. 2018. 46 (Issue D1): D348-D353.

[9] Gupta S, Stamatoyannopolous JA, Bailey T, Noble WS. *Quantifying similarity between motifs.* Genome Biology. 2007. 8(2):R24.

[10] Friedman, J., Hastie, T., Tibshirani, R. *Regularization paths for generalized linear models via coordinate descent.* Journal of Statistical Software 2010, 33(1):1.

[11] Zou, H., Hastie, T., Tibshirani, R. et al. *On the degrees of freedom of the lasso.* The Annals of Statistics. 2007; 35(5): 2173-2192.
